# Functional reconstitution of a bacterial CO_2_ concentrating mechanism in *E. coli*

**DOI:** 10.1101/2020.05.27.119784

**Authors:** Avi I. Flamholz, Eli Dugan, Cecilia Blikstad, Shmuel Gleizer, Roee Ben-Nissan, Shira Amram, Niv Antonovsky, Sumedha Ravishankar, Elad Noor, Arren Bar-Even, Ron Milo, David F. Savage

## Abstract

Many photosynthetic organisms employ a CO_2_ concentrating mechanism (CCM) to increase the rate of CO_2_ fixation via the Calvin cycle. CCMs catalyze ≈50% of global photosynthesis, yet it remains unclear which genes and proteins are required to produce this complex adaptation. We describe the construction of a functional CCM in a non-native host, achieved by expressing genes from an autotrophic bacterium in an engineered *E. coli* strain. Expression of 20 CCM genes enabled *E. coli* to grow by fixing CO_2_ from ambient air into biomass, with growth depending on CCM components. Bacterial CCMs are therefore genetically compact and readily transplanted, rationalizing their presence in diverse bacteria. Reconstitution enabled genetic experiments refining our understanding of the CCM, thereby laying the groundwork for deeper study and engineering of the cell biology supporting CO_2_ assimilation in diverse organisms.

**One Sentence Summary:** A bacterial CO_2_ concentrating mechanism enables *E. coli* to fix CO_2_ from ambient air.

## Introduction

Nearly all carbon in the biosphere enters by CO_2_ fixation in the Calvin-Benson-Bassham cycle (Raven et al., 2017). Ribulose Bisphosphate Carboxylase/Oxygenase - commonly known as rubisco - is the CO_2_ fixing enzyme in this cycle (Wildman, 2002) and likely the most abundant enzyme on Earth (Bar-On and Milo, 2019). As rubisco is abundant and central to biology, one might expect it to be an exceptional catalyst, but it is not. Photosynthetic rubiscos are modest enzymes, with carboxylation turnover numbers (*k*_*cat*_) ranging from 1-10 s^-1^ (Flamholz et al., 2019; Iñiguez et al., 2020). Moreover, all known rubiscos catalyze a competing oxygenation of the five-carbon organic substrate, ribulose 1,5-bisphosphate (Bowes and Ogren, 1972; Cleland et al., 1998; Flamholz et al., 2019).

Rubisco arose > 2.5 billion years ago, when Earth’s atmosphere contained little O_2_ and abundant CO_2_ (Fischer et al., 2016; Shih et al., 2016). In this environment, rubisco’s eponymous oxygenase activity could not have hindered carbon fixation or the growth of CO_2_-fixing organisms. Present-day atmosphere, however, poses a problem for plants and other autotrophs: their primary carbon source, CO_2_, is relatively scarce (≈0.04%) while a potent competing substrate, O_2_, is abundant (≈21%).

CO_2_ concentrating mechanisms (CCMs) arose multiple times over the last 2 billion years (Flamholz and Shih, 2020; Raven et al., 2017) and overcome this problem by concentrating CO_2_ near rubisco (Figure 1A). In elevated CO_2_ environments most active sites are occupied with CO_2_ and not O_2_. As such, high CO_2_ increases the rate of carboxylation and competitively inhibits oxygenation (Bowes and Ogren, 1972) thereby improving overall carbon assimilation (Figure 1B). Today, at least four varieties of CCMs are found in plants, algae and bacteria (Flamholz and Shih, 2020; Raven et al., 2017), organisms with CCMs are collectively responsible for ≈50% of global net photosynthesis (Raven et al., 2017), and some of the most productive human crops (e.g. maize and sugarcane) rely on CCMs.

**Figure 1.**
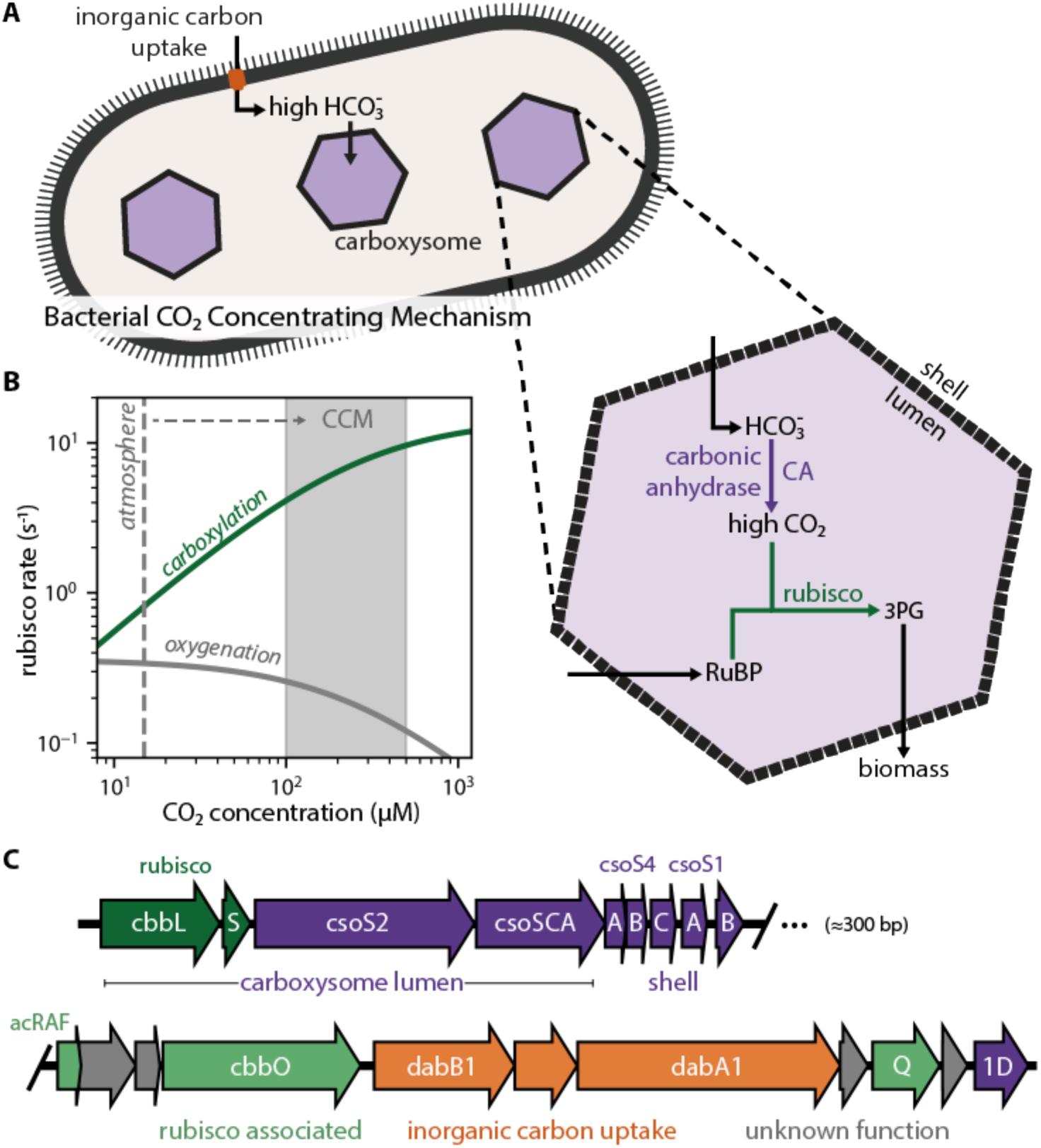
Twenty genes form the basis of a bacterial CCM. (**A**) The bacterial CCM consists of at least two essential components - energy-coupled carbon uptake and carboxysome structures that encapsulate rubisco with a carbonic anhydrase (CA) enzyme (Mangan et al., 2016; McGrath and Long, 2014). Transport generates a large cytosolic HCO_3_^-^ pool, which is rapidly converted to high carboxysomal CO_2_ concentration by the carboxysomal CA. (**B**) Elevated CO_2_ increases the rubisco carboxylation rate (green) and suppresses oxygenation by competitive inhibition (grey). [O_2_] was set to 270 μM for rate calculations. (**C**) H. neapolitanus CCM genes are mostly contained in a 20 gene cluster (Desmarais et al., 2019) expressing rubisco and its associated chaperones (green), carboxysome structural proteins (purple), and an inorganic carbon transporter (orange).

CCMs are particularly common among autotrophic bacteria: all Cyanobacteria and many Proteobacteria have CCM genes (Kerfeld and Melnicki, 2016; Rae et al., 2013). Bacterial CCMs rely on two crucial features: (i) energy-coupled inorganic carbon uptake at the cell membrane and (ii) a 200+ MDa protein organelle called the carboxysome that encapsulates rubisco with a carbonic anhydrase enzyme (Mangan et al., 2016; McGrath and Long, 2014). In the prevailing model of the carboxysome CCM, inorganic carbon uptake produces a high, above-equilibrium cytosolic HCO_3_^-^ concentration (≈30 mM) that diffuses into the carboxysome, where carbonic anhydrase activity produces a high carboxysomal CO_2_ concentration that promotes efficient carboxylation by rubisco (Figure 1A-B).

As CCMs accelerate CO_2_ fixation, there is great interest in transplanting them into crops (Ermakova et al., 2020; McGrath and Long, 2014). Carboxysome-based CCMs are especially attractive because they natively function in single cells and appear to rely on a tractable number of genes (Lin et al., 2014; Long et al., 2018; Occhialini et al., 2016; Orr et al., 2020). Modeling suggests that introducing bacterial CCM components could improve plant photosynthesis (McGrath and Long, 2014), especially if aspects of plant physiology can be modulated via genetic engineering (Wu et al., 2019). However, expressing bacterial rubiscos and carboxysome components has, so far, uniformly resulted in transgenic plants displaying impaired growth (Lin et al., 2014; Long et al., 2018; Occhialini et al., 2016; Orr et al., 2020). More generally, as our understanding of the genes and proteins participating in the carboxysome CCM rests mostly on loss-of-function genetic experiments in native hosts (Cai et al., 2009; Desmarais et al., 2019; Marcus et al., 1986; Price and Badger, 1989a), it is possible that some genetic, biochemical and physiological aspects of CCM function remain unappreciated. We therefore sought to test whether current understanding is sufficient to reconstitute the bacterial CCM in a non-native bacterial host, namely *E. coli*.

Using a genome-wide screen in the CO_2_-fixing proteobacterium *H. neapolitanus*, we recently demonstrated that a 20-gene cluster encodes all activities required for the CCM, at least in principle (Desmarais et al., 2019). These genes include rubisco large and small subunits, the carboxysomal carbonic anhydrase, seven structural proteins of the α-carboxysome (Bonacci et al., 2012), an energy-coupled inorganic carbon transporter (Desmarais et al., 2019; Scott et al., 2019), three rubisco chaperones (Aigner et al., 2017; Mueller-Cajar, 2017; Wheatley et al., 2014), and four genes of unknown function (Figure 1C). We aimed to test whether these genes are sufficient to establish a functioning CCM in *E. coli*.

## Results

As *E. coli* is a heterotroph, consuming organic carbon molecules to produce energy and biomass, it does not natively rely on rubisco. Therefore, in order to evaluate the effect of heterologous CCM expression, we first designed an *E. coli* strain that depends on rubisco carboxylation for growth. To grow on glycerol as the sole carbon source, *E. coli* must synthesize ribose 5-phosphate (Ri5P) for nucleic acids. Synthesis of Ri5P via the pentose phosphate pathway forces co-production of ribulose 5-phosphate (Ru5P). Deletion of ribose 5-phosphate isomerase (*rpiAB* genes, denoted Δrpi), however, makes Ru5P a metabolic “dead-end” (Figure 2A). Expression of phosphoribulokinase (*prk*) and rubisco creates a “detour” pathway converting Ru5P and CO_2_ into two units of the central metabolite 3-phosphoglycerate (3PG), enabling Ru5P metabolism and growth (Figure 2A). Additionally, cytosolic carbonic anhydrase activity is incompatible with the bacterial CCM (Price and Badger, 1989b). We therefore constructed a strain, named CCMB1 for “**CCM B**ackground **1**”, lacking *rpiAB* and all endogenous carbonic anhydrases (Methods).

**Figure 2.**
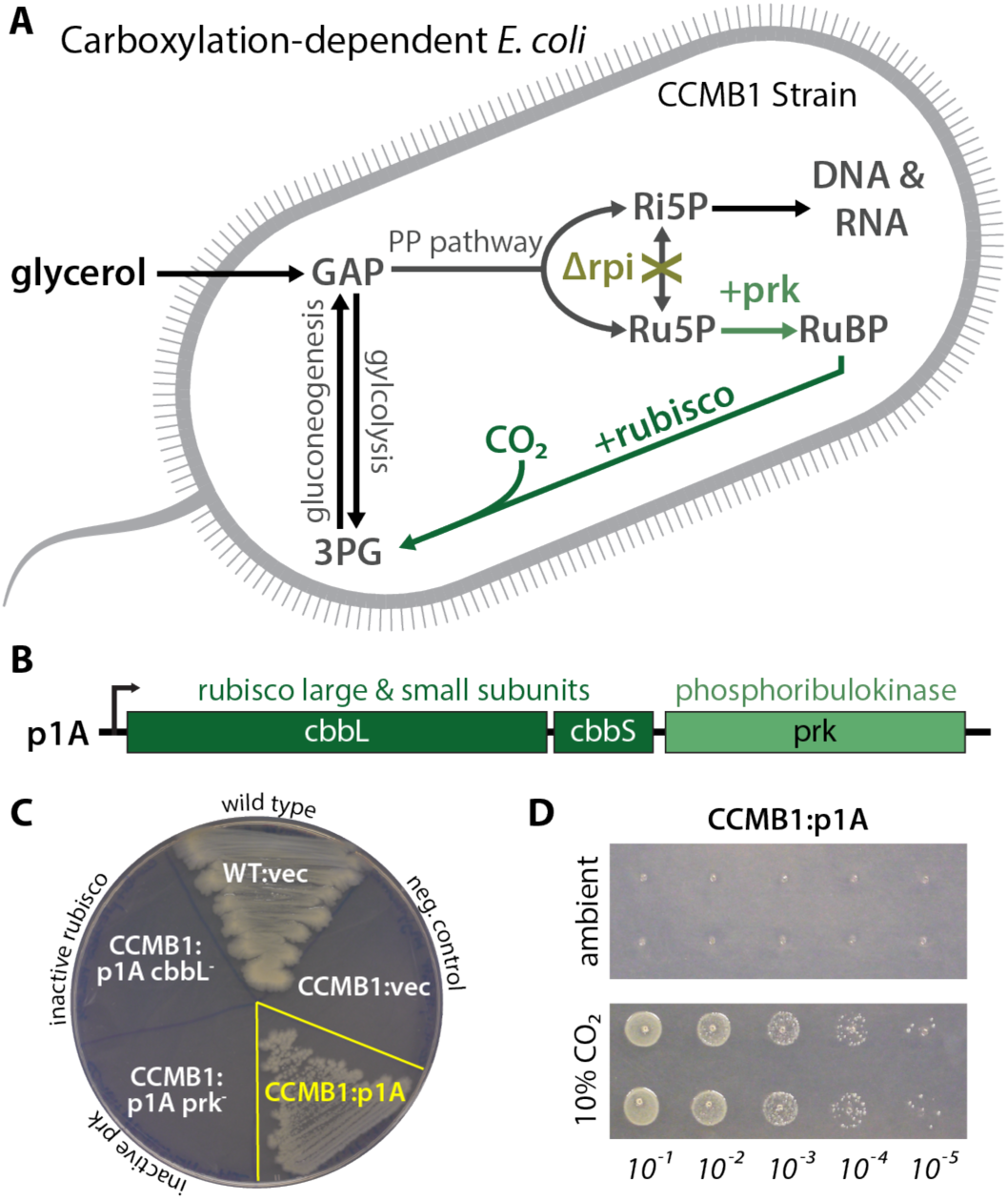
CCMB1 depends on rubisco carboxylation for growth on glycerol. (**A**) Ribose-5-phosphate (Ri5P) is required for nucleotide biosynthesis. Deletion of ribose-phosphate isomerase (Δrpi) in CCMB1 blocks ribulose-5-phosphate (Ru5P) metabolism in the pentose phosphate (PP) pathway. Expression of rubisco (H. neapolitanus cbbLS) and phosphoribulokinase (S. elongatus PCC7942 prk) on the p1A plasmid (**B**) permits Ru5P metabolism, thus enabling growth on M9 glycerol media in 10% CO_2_ (**C**). Mutating the rubisco active site (p1A cbbL^-^) abrogates growth, as does mutating ATP-binding residues of prk (p1A prk^-^). (**D**) CCMB1:p1A grows well under 10% CO_2_, but fails to grow in ambient air. Cells grown on M9 glycerol media throughout. The algorithmic design of CCMB1 is described in figure supplement 1 and the mechanism of rubisco-dependence is diagrammed in figure supplement 2. Figure supplement 3 shows CCMB1:p1A growth phenotypes on various media and figure supplement 4 demonstrates that rubisco oxygenation is not required for growth by demonstrating growth in the absence of O_2_. Acronyms: ribulose 1,5-bisphosphate (RuBP), 3-phosphoglycerate (3PG).

As predicted, CCMB1 required rubisco and prk for growth on glycerol minimal media in 10% CO_2_ (Figures 2B-C). When expressing rubisco and *prk* on the p1A plasmid (Figure 2B), CCMB1 also grew reproducibly in an anoxic mix of 10:90 CO_2_:N_2_ (Figure 2 - figure supplement 4) implying that rubisco carboxylation is sufficient for growth on glycerol media and rubisco-catalyzed oxygenation of RuBP is not required. CCMB1:p1A failed to grow on glycerol media in ambient air, however, presumably due to insufficient carboxylation at low CO_2_ (Figure 2D). That is, CCMB1:p1A displays the “high-CO_2_ requiring” phenotype that is the hallmark of CCM mutants (Marcus et al., 1986; Price and Badger, 1989a).

We expected that expressing a functional CO_2_-concentrating mechanism would cure CCMB1 of its high-CO_2_ requirement and permit growth in ambient air. We therefore generated two plasmids, pCB and pCCM, that together express all 20 genes from the *H. neapolitanus* CCM cluster (Figure 1C). pCB encodes ten carboxysome genes (Bonacci et al., 2012), including rubisco large and small subunits, along with *prk*. The remaining *H. neapolitanus* genes, including putative rubisco chaperones (Aigner et al., 2017; Mueller-Cajar, 2017; Wheatley et al., 2014) and an inorganic carbon transporter (Desmarais et al., 2019; Scott et al., 2019), were cloned into the second plasmid, pCCM.

CCMB1 co-transformed with pCB and pCCM initially failed to grow on glycerol media in ambient air. We therefore conducted selection experiments, described fully in Figure S5, that resulted in the isolation of mutant plasmids conferring growth in ambient air. Briefly, CCMB1:pCB + pCCM cultures were grown to saturation in 10% CO_2_. These cultures were washed and plated on glycerol minimal media (Methods). Colonies became visible after 20 days of incubation in ambient air (Figure S5). Deep-sequencing of plasmid DNA revealed mutations in regulatory sequences (e.g. a promoter and transcriptional repressor) but none in sequences coding for CCM components (Table S4). Individual post-selection plasmids pCB’ and pCCM’ were reconstructed by PCR, resequenced, and transformed into naive CCMB1 (Methods). As shown in Figure 3, pCB’ and pCCM’ together enabled reproducible growth of CCMB1 in ambient air, suggesting that the 20 genes expressed are sufficient to produce a heterologous CCM without any genomic mutations.

**Figure 3.**
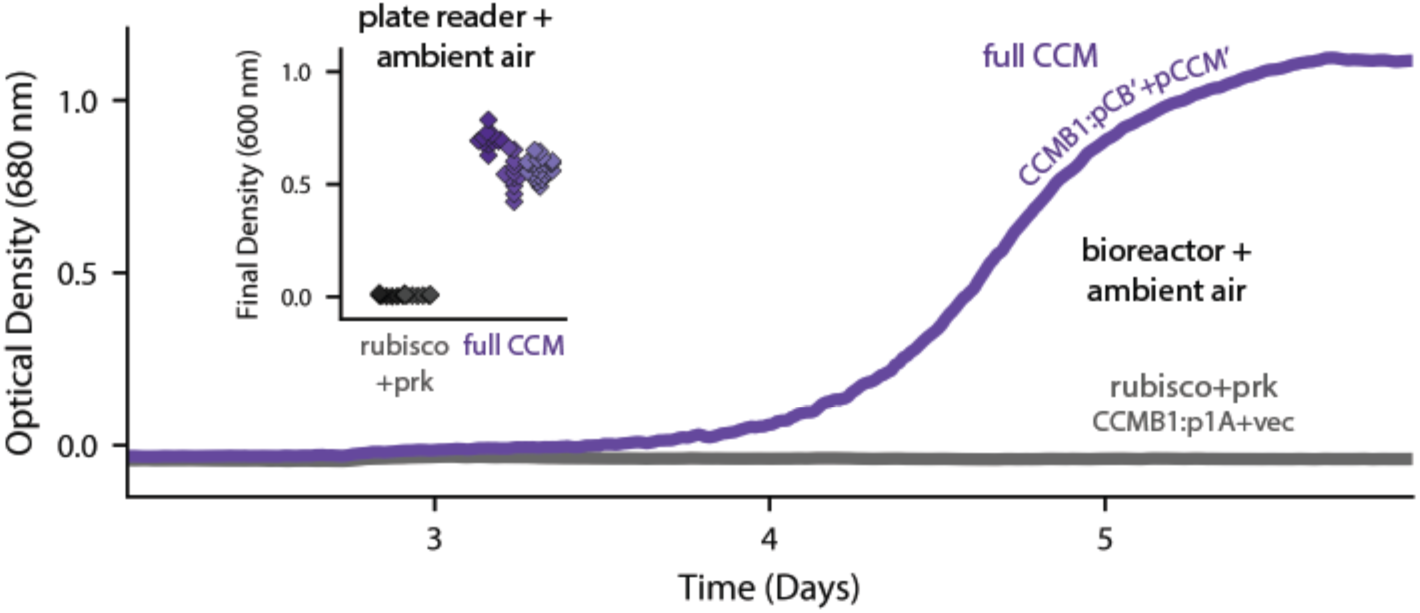
Expression of 20 CCM genes permits growth of CCMB1 in ambient air. Time course data give representative growth curves from a bioreactor bubbling ambient air. CCMB1:pCB’ + pCCM’ grows well (purple, “full CCM”), while rubisco and prk alone are insufficient for growth in ambient air (grey, CCMB1:p1A+vec). Inset: a plate reader experiment in biological triplicate (different shades) gave the same result. Expressing the full complement of CCM genes led to an increase in culture density (optical density at 600 nm) of ≈0.6 units after 80 hours of cultivation. Bootstrapping was used to calculate a 99.9% confidence interval of 0.56-0.64 OD units for the effect of expressing the full CCM during growth in ambient air. Figure supplement 1 shows triplicate growth curves and evaluates statistical significance.

To verify that growth in ambient air depends on the CCM, we generated plasmids carrying targeted mutations to known CCM components (Figure 4). An inactivating mutation to the carboxysomal rubisco (CbbL K194M) prohibited growth entirely. Mutations targeting the CCM, rather than rubisco itself, should ablate growth in ambient air while permitting growth in high CO_2_ (Desmarais et al., 2019; Mangan et al., 2016; Marcus et al., 1986; Price and Badger, 1989a; Rae et al., 2013). Consistent with this understanding, an inactive mutant of the carboxysomal carbonic anhydrase (CsoSCA C173S) required high-CO_2_ for growth. Similarly, disruption of carboxysome formation by removal of the pentameric shell proteins or the N-terminal domain of CsoS2 also eliminated growth in ambient air. Removing the pentameric proteins CsoS4AB disrupts the permeability barrier at the carboxysome shell (Cai et al., 2009), while truncating CsoS2 prohibits carboxysome formation entirely (Oltrogge et al., 2020). Finally, an inactivating mutation to the inorganic carbon transporter also eliminated growth in ambient air (Desmarais et al., 2019).

**Figure 4.**
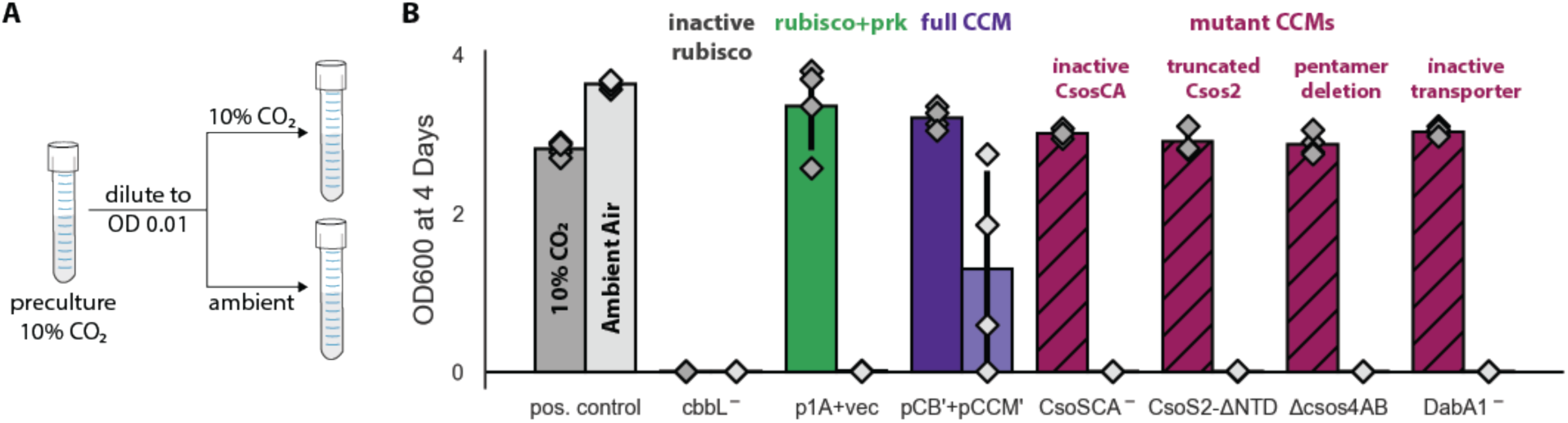
Growth in ambient air depends on the known components of the bacterial CCM. We generated plasmid variants carrying inactivating mutations to known components of the CCM. (**A**) Pre-cultures were grown in 10% CO_2_ and diluted into two tubes, one of which was cultured in 10% CO_2_ and the other in ambient air (Methods). Strains were tested in biological quadruplicate and culture density was measured after four days. (**B**) Targeted mutations to CCM components ablated growth in ambient air while permitting growth in 10% CO_2_, as expected. The left bar (darker color) gives the mean endpoint density in 10% CO_2_ for each strain. The right bar (lighter color) gives the mean in ambient air. Error bars give the standard deviation. From left to right: a positive control (grey) grew in 10% CO_2_ and ambient air, while a negative control CCMB1 strain carrying catalytically inactive rubisco (CCMB1:pCB’ cbbL^-^+pCCM’) failed to grow in either condition; CCMB1 expressing rubisco and prk but no CCM genes (green, CCMB1:p1A+vec) grew only in 10% CO_2_; CCMB1:pCB’+pCCM’ grew in 10% CO_2_ and ambient air, recapitulating results presented in Figure 3. The following four pairs of maroon bars give growth data for strains carrying targeted mutations to CCM genes: an inactivating mutation to carboxysomal carbonic anhydrase (CCMB1:pCB’ CsoSCA^-^+pCCM’), deletion of the CsoS2 N-terminus responsible for recruiting rubisco to the carboxysome (CCMB1:pCB’ CsoS2 ΔNTD +pCCM’), deletion of pentameric vertex proteins (CCMB1:pCB’ ΔcsoS4AB + pCCM’), and inactivating mutations to the DAB carbon uptake system (CCMB1:pCB’ DabA1^-^ + pCCM’). All four CCM mutations abrogated growth in air while permitting growth in 10% CO_2_. The positive control is the CAfree strain expressing human carbonic anhydrase II (Methods). Figure supplement 1 describes additional controls, statistical analyses, and a longer timescale replicate experiment (12 days) that additionally tests the contribution of rubisco chaperones to the CCM. Detailed description of all plasmid and mutation abbreviations is given in Table S2.

These experiments demonstrate that pCB’ and pCCM’ enable CCMB1 to grow in ambient air in a manner that depends on the known components of the bacterial CCM. To confirm that these cells produce carboxysome structures, we performed thin section electron microscopy. Regular polyhedral inclusions of ≈100 nm diameter were visible in micrographs (Figure 5A), implying production of morphologically-normal carboxysomes.

**Figure 5.**
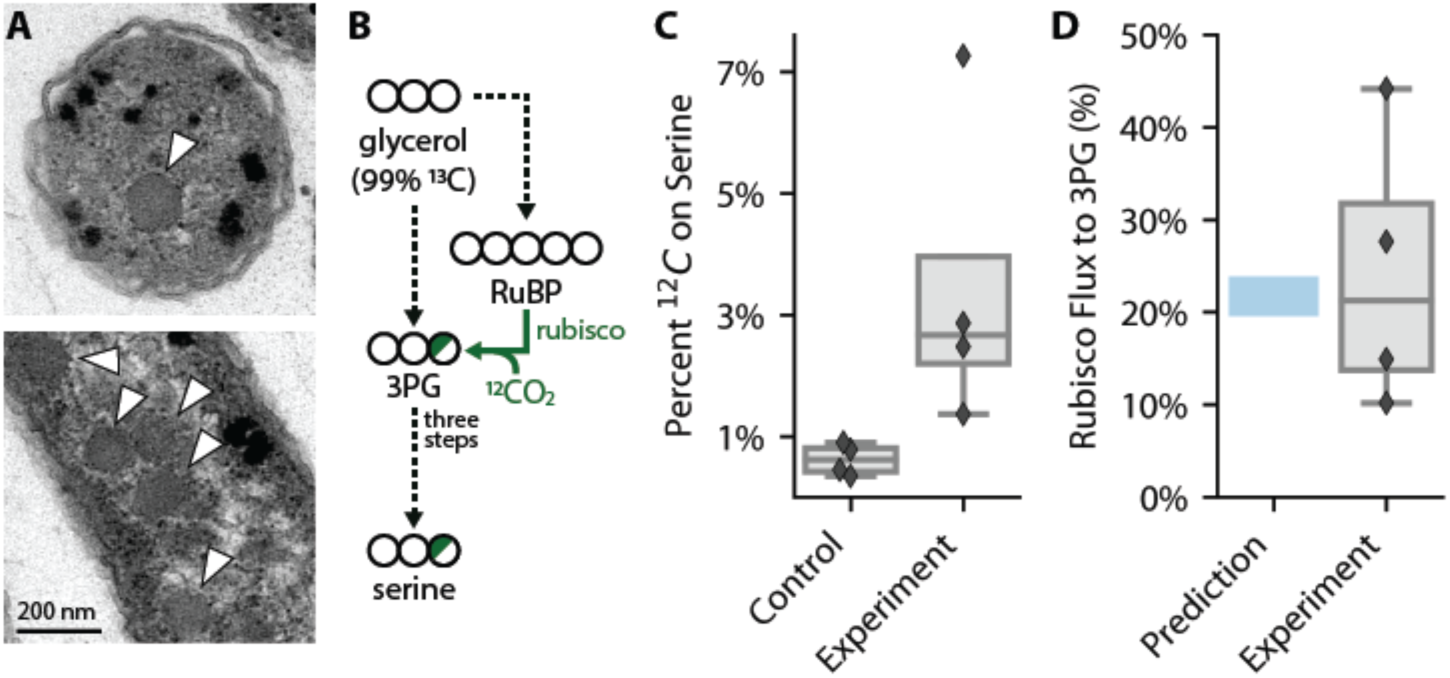
CCMB1:pCB’+pCCM’ produces morphologically normal carboxysomes and fixes CO_2_ from ambient air into biomass. (**A**) Polyhedral bodies resembling carboxysomes are evident in electron micrographs of CCMB1:pCB’+pCCM’ cells grown in air. Figure supplement 1 shows images of control strains. (**B**) Biological replicate cultures were grown in ambient air with 99% ^13^C glycerol as the sole organic carbon source so that ^12^CO_2_ in air is the sole source of ^12^C. As serine is a direct metabolic product of 3PG, we expect ^12^C enrichment on serine when rubisco is active. 3PG also derives from glycolytic metabolism of glycerol, so complete ^12^C labeling of serine was not expected. (**C**) The ^12^C composition of serine from CCMB1:pCB’ + pCCM’ (“Experiment”) is roughly threefold above the control. Figure supplement 2 gives ^12^C composition of all measured amino acids. (**D**) The fraction of 3PG production flux due to rubisco was predicted via Flux Balance Analysis and estimated from isotopic labeling data (Methods). Estimates of the rubisco flux fraction exceed 10% for all four biological replicates and the mean estimate accords well with a ≈20% prediction. Figure supplement 3 details the flux calculation procedure.

We next conducted isotopic labeling experiments to determine whether CCMB1:pCB’ + pCCM’ fixes CO_2_ from ambient air into biomass. Cells were grown in minimal media with ^13^C-labeled glycerol as the sole organic carbon source, such that CO_2_ from ambient air was the dominant source of ^12^C. The isotopic composition of amino acids in total biomass hydrolysate was analyzed via mass spectrometry (Methods). Serine is a useful sentinel of rubisco activity because *E. coli* produces serine from the rubisco product 3PG (Stauffer, 2004; Szyperski, 1995). 3PG is also an intermediate of lower glycolysis (Bar-Even et al., 2012), and so the degree of ^12^C labeling on serine reports on the balance of fluxes through rubisco and lower glycolysis (Figure 5B). We therefore expected excess ^12^C labeling of serine when rubisco is active. Consistent with this expectation, serine from CCMB1:pCB’+pCCM’ cells contained roughly threefold more ^12^C than the rubisco-independent control (Figure 5C). We estimated the contribution of rubisco to 3PG synthesis *in vivo* by comparing labeling patterns between the rubisco-dependent experimental cultures and controls (Methods). Based on these estimates, rubisco carboxylation was responsible for at least 10% of 3PG synthesis in all four biological replicates (Figure 5D, Methods), confirming fixation of CO_2_ from ambient air. As such, this work represents the first functional reconstitution of any CCM.

Reconstitution in *E. coli* enabled us to investigate which *H. neapolitanus* genes are necessary for CCM function in the absence of any regulation or genetic redundancy (i.e. genes with overlapping function) present in the native host. We focused on genes involved in rubisco proteostasis and generated plasmids lacking acRAF, a putative rubisco chaperone, or carrying targeted mutations to CbbQ, an ATPase involved in activating rubisco catalysis (Aigner et al., 2017; Mueller-Cajar, 2017; Wheatley et al., 2014). Although *acRAF* deletion had a large negative effect in *H. neapolitanus* (Desmarais et al., 2019), neither acRAF nor CbbQ were strictly required for CCMB1 to grow in ambient air. Consistent with our screen in the native host (Desmarais et al., 2019), however, *acRAF* deletion produced a substantial growth defect (Figure 4 - figure supplement 1, panel C), suggesting that the rate of rubisco complex assembly is an important determinant of carboxysome biogenesis.

## Discussion

Today, CCMs catalyze about half of global photosynthesis (Raven et al., 2017), but this was not always so. Land plant CCMs, for example, arose only in the last 100 million years (Flamholz and Shih, 2020; Raven et al., 2017; Sage et al., 2012). Though all contemporary Cyanobacteria have CCM genes, these CCMs are found in two convergently-evolved varieties (Flamholz and Shih, 2020; Kerfeld and Melnicki, 2016; Rae et al., 2013), suggesting that the ancestor of present-day Cyanobacteria and chloroplasts did not have a CCM (Rae et al., 2013). So how did carboxysome CCMs come to dominate the cyanobacterial phylum?

Here we demonstrated that the α-carboxysome CCM from *H. neapolitanus* is readily transferred between species and confers a large growth benefit, which can explain how these CCMs became so widespread among bacteria (Kerfeld and Melnicki, 2016; Rae et al., 2013). We constructed a CCM by expressing 20 genes in an engineered *E. coli* strain, CCMB1. In accordance with its role in native autotrophic hosts (Desmarais et al., 2019; Long et al., 2018; Marcus et al., 1986; Price and Badger, 1989a), the transplanted CCM required α-carboxysomes and inorganic carbon uptake to enable CCMB1 to grow by fixing CO_2_ from ambient air (Figures 3-5, S6-8). These results conclusively demonstrate that at most 20 gene products are required to produce a bacterial CCM. The α-carboxysome CCM is apparently genetically compact and “portable” between organisms. It is possible, therefore, that expressing bacterial CCMs in non-native autotrophic hosts will improve CO_2_ assimilation and growth. This is a promising approach to improving plant growth characteristics (Ermakova et al., 2020; Long et al., 2016; Wu et al., 2019) and also engineering enhanced microbial production of fuel, food products and commodity chemicals from CO_2_ (Claassens et al., 2016; Gleizer et al., 2019).

Reconstitution also enabled us to test, via simple genetic experiments, whether particular genes play a role in the CCM (Figure 4 - figure supplement 1). These experiments demonstrated that the rubisco chaperones are strictly dispensable for producing a functional bacterial CCM, though removing the *acRAF* gene produced a substantial growth defect that warrants further investigation. Further such can use our reconstituted CCM to delineate a minimal reconstitution of the bacterial CCM suitable for plant expression (Du et al., 2014; Long et al., 2018, 2016; Occhialini et al., 2016; Orr et al., 2020), test hypotheses about carboxysome biogenesis (Bonacci et al., 2012; Oltrogge et al., 2020), and probe the relationship between CCMs and host physiology (Mangan et al., 2016; Price and Badger, 1989b).

Our approach to studying CCMs by reconstitution in tractable non-native hosts can be applied to other CCMs, including β-carboxysome CCMs, the algal pyrenoid, and plausible evolutionary ancestors thereof (Flamholz and Shih, 2020). Historical trends in atmospheric CO_2_ likely promoted the evolution of CCMs (Fischer et al., 2016; Flamholz and Shih, 2020), so testing the growth of plausible ancestors of bacterial CCMs (e.g. carboxysomes lacking carbonic anhydrase activity) may provide insight into paths of CCM evolution and the composition of the ancient atmosphere at the time bacterial CCMs arose. In response to these same pressures, diverse eukaryotic algae evolved CCMs relying on micron-sized rubisco aggregates called the pyrenoids (Flamholz and Shih, 2020; Wang and Jonikas, 2020). Pyrenoid CCMs are collectively responsible for perhaps 80% of oceanic photosynthesis (Mackinder et al., 2016), yet many fundamental questions remain regarding the composition and operation of algal CCMs (Wang and Jonikas, 2020). Functional reconstitution of a pyrenoid CCM is a worthy goal which, once achieved, will indicate enormous progress in our collective understanding of the genetics, cell biology, biochemistry and physical processes supporting the eukaryotic complement of oceanic photosynthesis. We hope such studies will further our principled understanding of, and capacity to engineer, the cell biology supporting CO_2_ fixation in diverse organisms.

## Materials and Methods

### Growth conditions

Unless otherwise noted, cells were grown on M9 minimal media supplemented with 0.4% w/v glycerol, 0.5 ppm thiamin (10^4^ dilution of 0.5% w/v stock) and a trace element mix. The trace element mix components and their final concentrations in M9 media are: 50 mg/L EDTA, 31 mM FeCl_3_, 6.2 mM ZnCl_2_, 0.76 mM CuSO_4_·5H_2_O, 0.42 mM CoCl_2_·6H_2_O, 1.62 mM H_3_BO_3_, 81 nM MnCl_2_·4H_2_O. 100 nM anhydrotetracycline (aTc) was used in induced cultures. For routine cloning, 25 mg/L chloramphenicol and 60 mg/L kanamycin selection were used as appropriate. Antibiotics were reduced to half concentration (12.5 and 30 mg/L, respectively) for CCMB1 growth experiments and kanamycin was omitted when evaluating rubisco-dependence of growth as pF plasmids carrying kanamycin resistance also express rubisco. Culture densities were measured at 600 nm in a table top spectrophotometer (Genesys 20, Thermo Scientific) and turbid cultures were measured in five or tenfold dilution as appropriate in order to reach the linear regime of the spectrophotometer.

Agar plates were incubated at 37 °C in defined CO_2_ pressures in a CO_2_ controlled incubator (S41i, New Brunswick). For experiments in which a frozen bacterial stock was used to inoculate the culture, cells were first streaked on agar plates and incubated at 10% CO_2_ to facilitate fast growth. Pre-cultures derived from colonies were grown in 2-5 mL liquid M9 glycerol media under 10% CO_2_ with a matching 1 mL control in ambient air. Negative control strains unable to grow in minimal media (i.e. active site mutants of rubisco) were streaked on and pre-cultured in LB media under 10% CO_2_.

Growth curves were obtained using two complementary methods: an 8-chamber bioreactor for large-volume cultivation (MC1000, PSI), and 96-well plates in a gas controlled plate reader plate (Spark, Tecan). For the 96-well format, cells were pre-cultured in the appropriate permissive media, M9 glycerol under 10% CO_2_ where possible. If rich media was used, e.g. for negative controls, stationary phase cells were washed in 2x the culture volume and resuspended in 1x culture volume of M9 with no carbon source. Cultures were diluted to an OD of 1.0 (600 nm) and 250 μl cultures were inoculated by adding 5 μl of cells to 245 μl media. A humidity cassette (Tecan) was refilled daily with distilled water to mitigate evaporation during multi-day cultivation at 37 °C. Evaporation nonetheless produced irregular growth curves (e.g. Figure 2 - figure supplement 3), which motivated larger volume cultivation in the bioreactor, which mixes by bubbling ambient air into each growth vessel. 80 ml bioreactor cultures were inoculated to a starting OD of 0.005 (600 nm) and grown at 37 °C to saturation. Optical density was monitored continuously at 680 nm.

Anaerobic cultivation of agar plates was accomplished using a BBL GasPak 150 jar (BD) flushed 6 times with an anoxic mix of 10% CO_2_ and 90% N_2_. Tenfold titers of biological duplicate cultures were plated on M9 glycerol media with and without 20 mM NaNO_3_ supplementation. Because *E. coli* cannot ferment glycerol, NO_3_^-^ was supplied as an alternative electron acceptor. Plates without NO_3_^-^ showed no growth (Figure 2 - figure supplement 4), confirming the presence of an anaerobic atmosphere in the GasPak.

### Computational design of rubisco-dependent strains

To computationally design mutant strains in which growth is coupled to rubisco carboxylation flux, we used a variant of Flux Balance Analysis (Lewis et al., 2012) called “OptSlope” (Antonovsky et al., 2016). Starting from a published model of *E. coli* central metabolism, the Core *Escherichia coli* Metabolic Model (Orth et al., 2010), we considered all pairs of central metabolic knockouts and ignored those that permit growth *in silico* in the absence of rubisco and phosphoribulokinase (Prk) activities. For the remaining knockouts, we evaluated the degree of coupling between rubisco flux and biomass production during growth in nine carbon sources: glucose, fructose, gluconate, ribose, succinate, xylose, glycerate, acetate and glycerol. This approach highlighted several candidate rubisco-dependent knockout strains, including Δ*rpiAB* Δ*edd*. OptSlope predicted rubisco-dependent growth of Δ*rpiAB* Δ*edd* strains on glucose, fructose, succinate, acetate, glycerate, xylose and gluconate. The OptSlope algorithm is outlined in Figure 2 - figure supplement 1 and described fully in (Antonovsky et al., 2016). Proposed mechanisms of rubisco-dependence are outlined in Figure 2 - figure supplement 2. OptSlope source code is available at https://gitlab.com/elad.noor/optslope and calculations specific to CCMB1 can be found at https://github.com/flamholz/carboxecoli.

### Genomic modifications producing the CCMB1 strain

Strains used in this study are documented in Table S1. To produce CCMB1, we first constructed a strain termed “Δ*rpiAB*” for short. This strain has the genotype Δ*rpiAB* Δ*edd* and was constructed in the *E. coli* BW25113 background by repeated rounds of P1 transduction from the KEIO collection followed by pCP20 curing of the kanamaycin selection marker (Baba et al., 2006; Datsenko and Wanner, 2000). Deletion of *edd* removes the Entner-Doudoroff pathway (Peekhaus and Conway, 1998), forcing rubisco-dependent metabolism of gluconate via the pentose phosphate pathway (Figure 2 - figure supplement 2). CCMB1 has the genotype BW25113 Δ*rpiAB* Δ*edd* Δc*ynT* Δ*can* and was constructed from Δ*rpiAB* by deleting both native carbonic anhydrases using the same methods, first transducing the KEIO Δ*cynT* and then Δ*can* from EDCM636 (Merlin and Masters, 2003), which was obtained from the Yale Coli Genetic Stock Center. Transformation was performed by electroporation (ECM 630, Harvard Biosciences) and electrocompetent stocks were prepared using standard protocols. Strain genotypes were verified by PCR, as described below.

Plants, cyanobacteria and other autotrophs uniformly express “photorespiratory” pathways to process the rubisco oxygenation product 2-phosphoglycolate, or 2PG (Eisenhut et al., 2008). The *E. coli* genome encodes enzymes that could plausibly serve as a photorespiratory pathway (Figure 2 - figure supplement 2). We attempted to delete the *gph* gene in CCMB1 as it encodes the 2PG phosphatase that catalyzes the first step of this putative pathway. However, the Δ*gph* knockout was challenging to transform by electroporation, consistent with a proposed role in DNA repair (Pellicer et al., 2003). We reasoned that photorespiration might be required in CCMB1, as photorespiratory genes are essential in cyanobacteria (Eisenhut et al., 2008) and chemolithoautotrophic bacteria (Desmarais et al., 2019) even though both employ CCMs.

### Recombinant expression of rubisco, prk, and CCM components

pFE21 and pFA31 are compatible vectors derived from pZE21 and pZA31 (Lutz and Bujard, 1997). These vectors use an anhydrotetracycline (aTc) inducible P_LtetO-1_ promoter to regulate gene expression. pF plasmids were modified from parent vectors to constitutively express the tet repressor (TetR) under the P_bla_ promoter so that expression is repressed by default (Liang et al., 1999). We found that an inducible system aids in cloning problematic genes like *prk* (Wilson et al., 2018). We refer to these vectors as pFE and pFA respectively. The p1A plasmid (Figure 2A) derives from pFE and expresses two additional genes: the Form IA rubisco from *H. neapolitanus* and a *prk* gene from *Synechococcus elongatus PCC 7942*. The pCB plasmid is properly called pFE-CB, while pCCM is pFA-CCM. The two CCM plasmids are diagrammed in Figure 3 - figure supplement 1. Cloning was performed by Gibson and Golden-Gate approaches as appropriate. Large plasmids (e.g. pCB, pCCM) were verified by Illumina resequencing (Harvard MGH DNA Core plasmid sequencing service) and maps were updated manually after reviewing results compiled by breseq resequencing software (Deatherage and Barrick, 2014). Plasmids used in this study are described in Table S2 and available on Addgene at https://www.addgene.org/David_Savage/.

### Verifying the dependence of CCMB1 on rubisco carboxylation

To verify the dependence of CCMB1 on rubisco and Prk activities in minimal media, we constructed the variants of p1A carrying inactive rubisco or *prk* genes. Rubisco was inactivated by mutating the large subunit active site lysine to methionine, producing p1A CbbL K194M, or p1A CbbL^-^ for short (Andersson et al., 1989; Cleland et al., 1998). Prk was inactivated by mutating ATP-binding residues in the Walker A motif, producing p1A Prk K20M S21A, termed p1A Prk^-^ for short (Cai et al., 2014; Higgins et al., 1986). CCMB1:p1A grew on glycerol and gluconate minimal media when provided 10% CO_2_ (Figure 2 - figure supplement 3). CCMB1:p1A CbbL^-^ and CCMB1:p1A Prk^-^ both failed to grow on minimal media supplemented with glycerol or gluconate, demonstrating a dependence on both enzymes. So long as high CO_2_ was provided, neither activity was required for growth in rich LB media, which contains abundant nucleic acids precursors (Sezonov et al., 2007). Xylose minimal media was also tested but growth was impractically slow (data not shown).

The high-CO_2_ requirement of CCMB1:p1A growth was expected for two reasons: (i) bacterial rubiscos typically display low net carboxylation rates in ambient air due to relatively low CO_2_ (≈0.04%) and relatively high O_2_ (≈21%), as shown in Figure 1B and discussed in (Flamholz et al., 2019; Iñiguez et al., 2020), and (ii) CCMB1 entirely lacks carbonic anhydrase activity (Δ*cynT* Δ*can*). Carbonic anhydrase knockouts of many microbes, including *E. coli* and *S. cerevisiae*, are high-CO_2_ requiring, likely due to cellular demand for HCO_3_^-^ (Aguilera et al., 2005; Desmarais et al., 2019; Du et al., 2014; Merlin and Masters, 2003).

To verify that CCMB1 growth depends specifically on rubisco carboxylation and not oxygenation, we grew CCMB1:p1A on glycerol minimal medium in anoxic high-CO_2_ conditions (10:90 CO_2_:N_2_, Figure 2 - figure supplement 4). *E. coli* predominantly respires glycerol and, therefore, grows extremely slowly on glycerol in anaerobic and low O_2_ conditions (Stolper et al., 2010). We therefore supplied 20 mM NO_3_^-^ as an alternate terminal electron acceptor (Unden and Dünnwald, 2008) in anaerobic growth conditions (see “Growth conditions”). CCMB1:p1A grew on glycerol media in anaerobic conditions when NO_3_^-^ was provided. Growth is qualitatively weaker than a wild-type control, but this is consistent with the growth differences observed in aerobic conditions (Figure 2 - figure supplement 4). Anaerobic growth of CCMB1:p1A on glycerol minimal media implies that growth can be supported by rubisco carboxylation alone and does not require the rubisco-catalyzed oxygenation of RuBP.

### Strain verification by PCR and phenotypic testing

As CCMB1 is a relatively slow-growing knockout strain, we occasionally observed contaminants in growth experiments. We used two strategies to detect contamination by faster-growing organisms (e.g. wild-type *E. coli*). As most strains grew poorly or not at all in ambient air, pre-cultures grown in 10% CO_2_ were accompanied by a matching 1 mL negative control in ambient air. Pre-cultures showing growth in the negative control were discarded or verified by PCR genotyping in cases where air-growth was plausible.

PCR genotyping was performed using primer sets documented in Table S3. Three primer pairs were used to probe a control locus (*zwf*) and two target loci (*cynT* and *rpiA*). The *zwf* locus is intact in all strains. *cynT* and *rpiA* probes test for the presence of the CCMB1 strain (genotype BW25113 Δ*rpiAB* Δ*edd* Δ*cynT* Δ*can*). Notably, the CAfree strain (BW25113 Δ*cynT* Δ*can*) that we previously used to test the activity of DAB-type transporters (Desmarais et al., 2019) is a *cynT* knockout but has a wild-type *rpiA* locus, so this primer set can distinguish between wild-type, CAfree and CCMB1. This was useful for some experiments where CAfree was used as a control (e.g. Figures S7-8). Pooled colony PCRs were performed using Q5 polymerase (NEB), annealing at 65 °C and with a 50 second extension time.

### Selection for growth in novel conditions

CCMB1:pCB did not initially grow in glycerol minimal media, which was unexpected because pCB carries rubisco and *prk* genes. We therefore performed a series of selection experiments (Herz et al., 2017) to isolate plasmids conferring growth at elevated CO_2_ and then in ambient air. We first describe the methodology; the full series of experiments is diagrammed fully in Figure S5B and described in paragraphs below. CCMB1 cultures carrying appropriate plasmids were first grown to saturation in rich LB media in a 10% CO_2_ incubator. Stationary phase cultures were pelleted by centrifugation for 10 min at 4,000 x g, washed in 2x the culture volume, and resuspended in 1x culture volume of M9 media with no carbon source. After resuspension, multiple dilutions were plated on selective media (e.g. M9 glycerol media) and incubated in the desired conditions (e.g. in ambient air) with a positive control in 10% CO_2_ on appropriate media. In later experiments, matching tenfold titers were plated in permissive conditions (e.g. in 10% CO_2_) to estimate the number of viable cells. When colonies formed in restrictive conditions, they were picked into permissive media, grown to saturation, washed and tested for re-growth in restrictive conditions by titer plating or streaking. Plasmid DNA was isolated from verified colonies and transformed into naive CCMB1 cells to test whether plasmid mutations confer improved growth (i.e. in the absence of genomic mutations).

We first selected for CCMB1:pCB growth on M9 glycerol media in 10% CO_2_ and then in M9 gluconate media under 10% CO_2_. Pre-cultures to stationary phase in rich media in 10% CO_2_, and then plated on selective media after washing. The resulting plasmid, pCB-gg for “gluconate grower,” was isolated and deep sequenced (Harvard MGH DNA Core plasmid sequencing service). Plasmid maps were updated manually after running the breseq resequencing software (Deatherage and Barrick, 2014). pCB-gg was found to carry two regulatory mutations: an amino acid substitution to the tet repressor (TetR E37A) and a nucleotide substitution in the Tet operator regulating the carboxysome operon (tetO_2_ +8T, Table S4).

Following this first round of selection, CCMB1 was co-transformed with pCB-gg and pCCM. The transformants grew in M9 glycerol media in 10% CO_2_ but failed to grow on in ambient air. We therefore performed another selection experiment, plating CCMB1:pCB-gg+pCCM on M9 glycerol media in ambient CO_2_. Parallel negative control selections were conducted on uninduced plates (no aTc) and using CCMB1:p1A+pCCM, which lacks carboxysome genes. Colonies formed on induced CCMB1:pCB-gg+pCCM plates after 20 days, but not on control plates (Figure S5F).

Forty colonies were picked and tested for re-growth in ambient CO_2_ by tenfold titer plating. 10 of 40 regrew (six examples are shown in Figure S5G). Pooled plasmid DNA was extracted from verified colonies and electroporated into naive CCMB1 to test plasmid-linkage of growth. We found that plasmid DNA from colony #4 produced the most robust growth in ambient air. This was tested by picking 16 re-transformants and testing their growth in ambient air in liquid M9 glycerol media. Re-transformant #13 regrew robustly in all 6 technical replicates. Pooled plasmid DNA from colony #4 re-transformant #13 was resequenced by a combination deep sequencing (as above) and targeted Sanger sequencing of the TetR locus and origins of replication, as these regions share sequence between both parent plasmids. pCB carried the same mutations as pCB-gg and pCCM had acquired the high-copy ColE1 origin of replication from pCB (Table S4). The individual mutant plasmids, termed pCB’ and pCCM’, were reconstructed from pooled plasmid extract by PCR and Gibson cloning.

These post-selection plasmids, termed pCB’ and pCCM’, were again verified by resequencing. Naive CCMB1 was transformed with the reconstructed post-selection plasmids pCB’ and pCCM’ and tested for growth in ambient air. We found that the post-selection plasmids confer reproducible growth in ambient air in multiple growth conditions (Figure 3), implying that genomic mutations that formed during selections were not required to produce growth in ambient air.

### Design of mutant CCM plasmids

To verify that air-growth depends on the known components of the CCM, we generated variants of pCB’ and pCCM’ carrying known, targeted null mutations to the CCM. CCMB1 was co-transformed with two plasmids: a mutant plasmid (of either pCB’ or pCCM’) and its cognate, unmodified plasmid. Mutant plasmids are listed here along with expected growth phenotypes, with fuller detail in Table S2. pCB’ CbbL K194M, or pCB’^-^, contains an inactivating mutation to the large subunit of the carboxysomal Form 1A rubisco (Andersson et al., 1989; Cleland et al., 1998). This mutation was expected to abrogate rubisco-dependent growth entirely.

Mutations targeting the CCM, rather than rubisco itself, are expected to ablate growth in ambient air but permit growth in high CO_2_. The following plasmid mutations were designed to specifically target essential components of the CCM. pCB’ CsoSCA C173S, or pCB’ CsoSCA^-^, carries a mutation to an active site cysteine residue responsible for coordinating the catalytic Zn2+ ion in **β**-carbonic anhydrases (Sawaya et al., 2006). pCB’ CsoS2 ΔNTD lacks the N-terminal domain of CsoS2, which is responsible for recruiting rubisco to the carboxysome during the biogenesis of the organelle (Oltrogge et al., 2020). Similarly, pCB’ CbbL Y72R carries an arginine residue instead of the tyrosine responsible for mediating cation-π interactions between the rubisco large subunit and the N-termus of CsoS2. This mutation was shown to eliminate any binding interaction between the rubisco complex and the N-termus of CsoS2 (Oltrogge et al., 2020). pCB’ Δ*csoS4AB* lacks both pentameric shell proteins, CsoS4AB, which was shown to disrupt the permeability barrier at the carboxysome shell (Cai et al., 2009). pCCM’ DabA1 C462A, D464A, or pCCM’ DabA1^-^, carries inactivating mutations to the putative active site of the inorganic carbon transporter component DabA1 (Desmarais et al., 2019).

Two more mutant plasmids were designed to test the roles of rubisco chaperones in producing a functional CCM. pCCM’ CbbQ K46A, E107Q, denoted pCCM’ CbbQ^-^, carries mutations that inactivate the ATPase activity of the CbbQ subunit of the CbbOQ rubisco activase complex (Tsai et al., 2015). pCCM’ Δ*acRAF* lacks the putative rubisco chaperone acRAF. acRAF is homologous to a plant rubisco folding chaperone (Aigner et al., 2017) and likely involved in the folding of the *H. neapolitanus* Form IA rubisco (Wheatley et al., 2014). Experimental evaluation of growth phenotypes for the above-described mutants is detailed below and results are given in Figure 4 - figure supplement 1.

### Phenotyping of matched cultures in 10% CO_2_ and ambient air

To interrogate the phenotypic effects of mutations to the CCM, we tested the growth of matched biological replicate cultures of CCM mutants (e.g. disruption of carboxysome components or transporter function) in M9 glycerol medium in 10% CO_2_ and ambient air (Figure 4A). For these experiments, individual colonies were picked into a round-bottom tube with 4 mL of M9 glycerol media with full strength antibiotic and 100 nM aTc. 1 mL of culture was then transferred to a second tube. The 3 mL pre-culture was incubated in 10% CO_2_, while the 1 mL culture was incubated in ambient air as a negative control. Control strains unable to grow in minimal media (e.g. those expressing inactive rubisco mutants) were pre-cultured in LB media. High-CO_2_ pre-cultures were grown to saturation, after which optical density (OD600) was measured in five-fold dilution.

Experimental cultures were inoculated with pre-culture to a starting OD600 of 0.01 in 3 mL of M9 glycerol media with 12.5 mg/L chloramphenicol and 100 nM aTc. Each pre-culture was used to inoculate a matched pair of experimental cultures, one incubated in 10% CO_2_ and another in ambient air, as diagrammed in Figure 4A. After a defined period of growth (4 days in Figure 4B and 12 days in Figure 4 - figure supplement 1, panel C) all culture densities were measured at 600 nm. All experiments were performed in biological quadruplicate, i.e. using four independent pre-cultures deriving from distinct colonies to inoculate four pairs of matched cultures. A positive control was included in all experiments to test the media composition. We used a complemented double carbonic anhydrase knockout (CAfree:pFE-sfGFP+pFA-HCAII) for this purpose as its growth in air depends on the expression of the human carbonic anhydrase II from the pFA-HCAII plasmid (Desmarais et al., 2019).

### Electron microscopy

CCMB1:pCB’+pCCM’ was grown in ambient air in 3 ml of M9 glycerol medium and induced with 100 nM aTc. A carboxysome-negative control, CAfree:pFE-sfGFP+pFA-HCAII, was grown in the same conditions. Sample preparation and sectioning were performed by the University of California Berkeley Electron Microscope Laboratory. Cell pellets were fixed for 30 min at room temperature in 2.5% glutaraldehyde in 0.1 M cacodylate buffer pH 7.4. Fixed cells were stabilized in 1% very low melting-point agarose and cut into small cubes. Cubed sample was then rinsed three times at room temperature for 10 min in 0.1 M sodium cacodylate buffer, pH 7.4 and then immersed in 1% osmium tetroxide with 1.6% potassium ferricyanide in 0.1 M cacodylate buffer for an hour in the dark on a rocker. Samples were later rinsed three times with a cacodylate buffer and then subjected to an ascending series of acetone for 10 min each (35%, 50%, 75%, 80%, 90%, 100%, 100%). Samples were progressively infiltrated with Epon resin (EMS, Hatfield, PA, USA) while rocking and later polymerized at 60 °C for 24 hours. 70 nm thin sections were cut using an Ultracut E (Leica) and collected on 100 mesh formvar coated copper grids. The grids were further stained for 5 min with 2% aqueous uranyl acetate and 4 min with Reynold’s lead citrate. The sections were imaged using a Tecnai 12 TEM at 120 KV (FEI) and images were collected using UltraScan 1000 digital micrograph software (Gatan Inc.).

### Sample preparation for LC-MS analysis

Protein-bound amino acids were analyzed in total biomass hydrolysate of 80 mL cultures grown in minimal media with 99% ^13^C glycerol (Cambridge Isotopes) as the sole organic carbon source. These cultures were grown in 80 mL volumes in a bioreactor pumping ambient air (MC1000, PSI). After harvesting biomass, samples were prepared and analyzed as described in (Antonovsky et al., 2016). Briefly, the OD600 was recorded and 0.6 OD x mL of sample were pelleted by centrifugation for 15 min at 4,000 x g. The pellet was resuspended in 1 mL of 6 N HCl and incubated for 24 hours at 110 °C. The acid was subsequently evaporated under a nitrogen stream using a custom-built gas manifold (Nevins et al., 2005), resulting in a dry hydrolysate. Dry hydrolysates were resuspended in 0.6 mL of MilliQ water, centrifuged for 5 min at 14,000 x g, and supernatant was analyzed by liquid chromatography-mass spectrometry (LC-MS). Hydrolyzed amino acids were separated using ultra performance liquid chromatography (UPLC, Acquity, Waters) on a C-8 column (Zorbax Eclipse XBD, Agilent) at a flow rate of 0.6 mL/min, and eluted off the column using a hydrophobicity gradient. Buffers used were: A) H2O + 0.1% formic acid and B) acetonitrile + 0.1% formic acid with the following gradient: 100% of A (0-3 min), 100% A to 100% B (3-9 min), 100% B (9-13 min), 100% B to 100% A (13-14 min), 100% A (14-20 min). The UPLC was coupled online to a triple quadrupole mass spectrometer (TQS, Waters). Data were acquired using MassLynx v4.1 (Waters). Amino acids and metabolites used for analysis were selected according to the following criteria: amino acids that had peaks at a distinct retention time and m/z values for all isotopologues and also showed correct ^13^C labeling fractions in control samples that contained protein hydrolyzates of WT cells grown with known ratios of ^13^C6-glucose to ^12^C-glucose.

### Isotopic analysis composition of biomolecules

The total ^13^C fraction of each metabolite was determined as the weighted average of the fractions of all the isotopologues for that metabolite: 

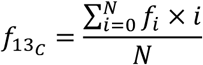

Here N is the number of carbons in the compound (e.g. N = 3 for serine) and *f*_*i*_ is the relative fraction of the i-th isotopologue, i.e containing *i* ^13^C carbon atoms. Each metabolite’s total ^12^C fraction was calculated as 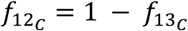.

### Estimating the effective intracellular ^12^CO_2_ fraction

*E. coli* cells grown in ^13^C glycerol will simultaneously respire glycerol, producing intracellular ^13^CO_2_, and take up extracellular ^12^CO_2_ and H^12^CO_3_^-^. The isotopic composition of the intracellular inorganic carbon (C_i_) pool will therefore reflect the balance of uptake and respiration. As rubisco carboxylation draws from the intracellular CO_2_ pool, we must estimate the isotopic composition of the C_i_ pool to evaluate the contribution of rubisco to metabolism. We used the carbamoyl-phosphate moiety as a marker for the isotopic distribution of the intracellular C_i_ pool, as described in (Gleizer et al., 2019). Briefly, carbamoyl-phosphate is generated by phosphorylation of bicarbonate, and is condensed with ornithine in the biosynthesis of L-arginine. We compared the mass isotopologue distribution of L-arginine, which contains one carbon from carbamoyl-phosphate, with the mass isotopologue distribution of L-glutamate as L-glutamate is an ornithine precursor.

We estimated the effective ^13^C labeling of intracellular inorganic carbon 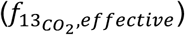, as follows: 

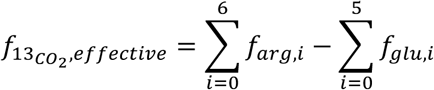

Here 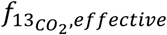 is the relative fraction of ^13^CO_2_ out of the total CO_2_ pool (or, more formally, the C_i_), and *f*_*arg,i*_ and *f*_*glu,i*_ are the fraction of the i-th isotopologue of arginine and glutamate respectively. We assumed fast equilibration of the intracellular C_i_ pool because the strains used in labeling experiments express a carbonic anhydrase. An equivalent equation can be defined for the arginine-proline comparison (Gleizer et al., 2019), however proline data were of insufficient quality to use and so we report inferences based on the arginine-glutamate comparison. The effective intracellular fraction of ^12^CO_2_ was calculated as 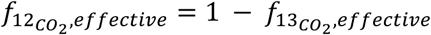. For brevity, we refer to these fractions as 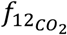 and 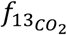, respectively.

### Estimating the rubisco carboxylation flux *in vivo*

When CCMB1 cells are grown on 99% ^13^C glycerol, 3-phosphoglycerate (3PG) can be produced via two routes: (i) rubisco catalyzed carboxylation of RuBP and (ii) glycolytic metabolism of glycerol via dihydroxyacetone phosphate, or DHAP (Booth, 2005). We denote these two fluxes as *J*_*rubisco*_ and *J*_*pgk*_, where pgk (phosphoglycerate kinase) is the glycolytic enzyme producing 3PG (Bar-Even et al., 2012). Serine is a direct metabolic product of 3PG (Stauffer, 2004; Szyperski, 1995) and was therefore assumed to have the same ^12^C composition as 3PG. Rubisco-catalyzed carboxylation of RuBP adds one CO_2_ to the 5-carbon substrate, producing two 3PG molecules containing a total of six carbon atoms. Therefore, ⅙ of carbon atoms on 3PG produced via rubisco carboxylation must derive from an inorganic source (Figure 5 - figure supplement 3). Carboxylation draws CO_2_ from the intracellular inorganic carbon pool, whose ^12^C composition 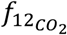 was inferred as described above.

Based on these assumptions, the ^12^C composition of 3PG, and therefore serine, equals a flux-weighted sum of contributions from rubisco and pgk. As such, the relative 3PG production flux that is due to rubisco, *J*_*rubisco*_*/(J*_*rubisco*_+*J*_*pgk*_), can be inferred via the following calculation: 

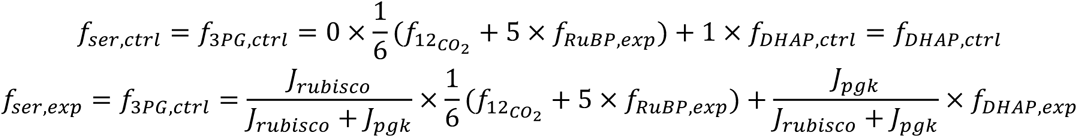

Where the first equation is written for the control and the second for experimental cultures where rubisco is active (CCMB1:pCB’+pCCM’). *f*_*ser,ctrl*_ and *f*_*ser,exp*_ denote the ^12^C composition of serine in the control and experiment respectively. Identical notation is used for RuBP and DHAP. As there are only two routes of 3PG production, the above equations can be simplified to solve for the relative flux through rubisco *in vivo*: 

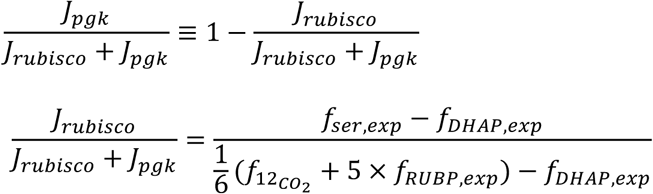

To calculate the rubisco flux *in vivo* we must attach values to several parameters in the above equation. 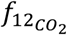 was inferred on a per-sample basis, with the mean values being 25% ± 4%and 67% ± 28% for the control and experiment respectively (Figure 5 - figure supplement 3, panel C). Because glycerol is converted into 3PG and serine via DHAP in wild-type *E. coli* (Booth, 2005), we expect that *f*_*ser,ctrl*_ = *f*_*DHAP,ctrl*_, as derived above. LC-MS measurements give *f*_*ser,ctrl*_ = 0.6% ± 0.2% and *f*_*ser,exp*_ = 3.5% ± 2.2%(Figure 5C). Valine is also a metabolic product of DHAP (Szyperski, 1995) and was found to have a similar ^12^C fraction *f*_*val,ctrl*_ = 0.6% ± 0.1% in control cells (Figure 5 - figure supplement 2). Since glycerol is immediately converted to DHAP in *E. coli*, we further assumed that *f*_*DHAP,ctrl*_ = *f*_*DHAP,exp*_.

RuBP is produced in CCMB1 when rubisco and prk are expressed. Since glycerol is the sole carbon source and there are no carboxylation reactions between DHAP and RuBP in CCMB1, we assumed *f*_*RuBP,exp*_ = *f*_*DHAP,ctrl*_. This assumption is supported by LC-MS measurements of histidine in control cells. Like RuBP, histidine is synthesized from a pentose-phosphate pathway intermediates (Szyperski, 1995; Winkler and Ramos-Montañez, 2009), and measured *f*_*his,ctrl*_ = 0.7% ± 0.1%, which is very similar to *f*_*ser,ctrl*_ = 0.6% ± 0.3%. Using mean values to illustrate the calculation gives 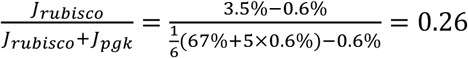, implying that 26% of 3PG production is due to rubisco.

10^5^ random samples were drawn from the experimentally determined parameter ranges to estimate a 99% confidence interval on the rubisco flux fraction. As the ^12^C composition of inorganic carbon 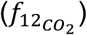 and serine are mechanistically linked via rubisco, these values were assumed to co-vary. Distributions were estimated on a per-sample basis by assuming 0.1% error in direct measurement of serine and 1% error in the inference of 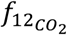. These calculations gave a median flux estimate of 19% with 99% of values falling between 4.0% and 47.3%. The sample with the lowest inferred rubisco flux had a median estimate of 10.2% with 99% of values falling between 2.5% and 17.9%, implying that rubisco is responsible for a nonzero fraction of 3PG production in all samples. Applying a wider error range of 5% to 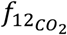 did not qualitatively change results, giving an overall median flux fraction of 19.1% and a 99% confidence interval 3.9-50.7%. This and above calculations can be found in the following Jupyter notebook: https://github.com/flamholz/carboxecoli/blob/master/notebooks/00_LCMS_calcs.ipynb.

### Predicting rubisco carboxylation flux via Flux Balance Analysis

A stoichiometric model of complemented CCMB1 was generated from the Core *Escherichia coli* Metabolic Model (Orth et al., 2010) by adding rubisco and prk and then deleting the rpi and edd reactions. Parsimonious Flux Balance Analysis (pFBA) was applied to the resulting model to calculate intracellular metabolic metabolic fluxes that maximize the rate of biomass production. As many distinct flux distributions can yield the same (maximal) rate of biomass production, pFBA uses the minimum sum of fluxes objective to define a unique flux solution (Holzhütter, 2004). The COBRApy implementation of pFBA introduces an additional free parameter, the permissible fraction of the maximal biomass production rate *f*_*opt*_ (Ebrahim et al., 2013). When *f*_*opt*_ < 1.0, the biomass production can be less-than-optimal if this would further decrease the sum of fluxes.

pFBA was run with *f*_*opt*_ ranging from 0.8 to 1.0 in increments of 0.01 to account for the fact that CCMB1 has not undergone selection to maximize biomass production with rubisco expressed. For each resulting flux distribution the fraction of 3PG production flux due to rubisco was calculated as the fraction of 3PG molecules produced via rubisco carboxylation divided by the total flux to 3PG. These calculations predict that 19.5%-21.5% of 3PG production is due to rubisco. The model was rerun after removing all possibility for product secretion by deleting all carbon-containing exchange reactions other than glycerol and CO_2_ exchange. This modification should give an upper bound on the fraction of 3PG production due to rubisco, as carbon cannot be shunted away from biomass production to overflow products. The “no overflow” model predicted that 23.9% of 3PG production is due to rubisco independent of *f*_*opt*_. The range of predictions from 19.5-23.9% is plotted in Figure 5D. All calculations were done using Python and COBRApy (Ebrahim et al., 2013), and can be found in this Jupyter notebook: https://github.com/flamholz/carboxecoli/blob/master/notebooks/01_FBA_rubisco_flux_prediction.ipynb.

## Acknowledgements

We thank Matt Davis for P1 transduction materials and advice, Hernan Garcia and Han Lim for pZ plasmids, Maggie Stoeva, Anna Engelbrektson, Anchal Mehra, Sophia Ewens and Tyler Barnum for help with anaerobic growth, Reena Zalpuri and Danielle Jorgens at the University of California Berkeley Electron Microscope Laboratory for advice and assistance with electron microscopy, and Rob Egbert and Adam Arkin for KEIO strains. We are grateful to Eric Estrin, Woody Fischer, Darcy McRose, Dipti Nayak, Sabeeha Merchant, Luke Oltrogge, and Naiya Phillips for detailed comments on the manuscript, and to Dan Arlow, Yinon Bar-On, Dan Davidi, Jack Desmarais, Hernan Garcia, Oliver Mueller-Cajar, Rob Nichols, Kris Niyogi, Dan Portnoy, Morgan Price, Noam Prywes, Jeremy Roop, Rachel Shipps, Patrick Shih, and Dan Tawfik, for support, advice and helpful discussions throughout.

## Author contributions

A.I.F. conceived of and designed all experiments with mentorship from R.M. and D.F.S. and support from all authors. S.A., N.A., E.N., A.B-E. and R.M. designed and constructed the Δ*rpiAB* strain from which A.I.F., E.J.D, and S.R. constructed CCMB1. A.I.F., E.J.D, and S.R. designed and constructed all other strains and plasmids. A.I.F and E.J.D. performed growth and selection experiments. C.B. performed electron microscopy. S.G. and R.B-N. performed LC-MS analysis on biomass hydrolysate prepared by A.I.F. and E.J.D. A.I.F., S.G., R.B-N., and E.N. analysed isotopic labeling data. A.I.F and E.N. designed and executed Flux Balance Analysis. A.I.F. wrote the manuscript with input from all authors.

## Data and materials availability

All data associated with this work is available in the main text, supplementary materials and the repository at https://github.com/flamholz/carboxecoli. Plasmids available on Addgene at https://www.addgene.org/David_Savage/, strains distributed on request.

## Figures Supplements

**Figure 2 - figure supplement 1.**
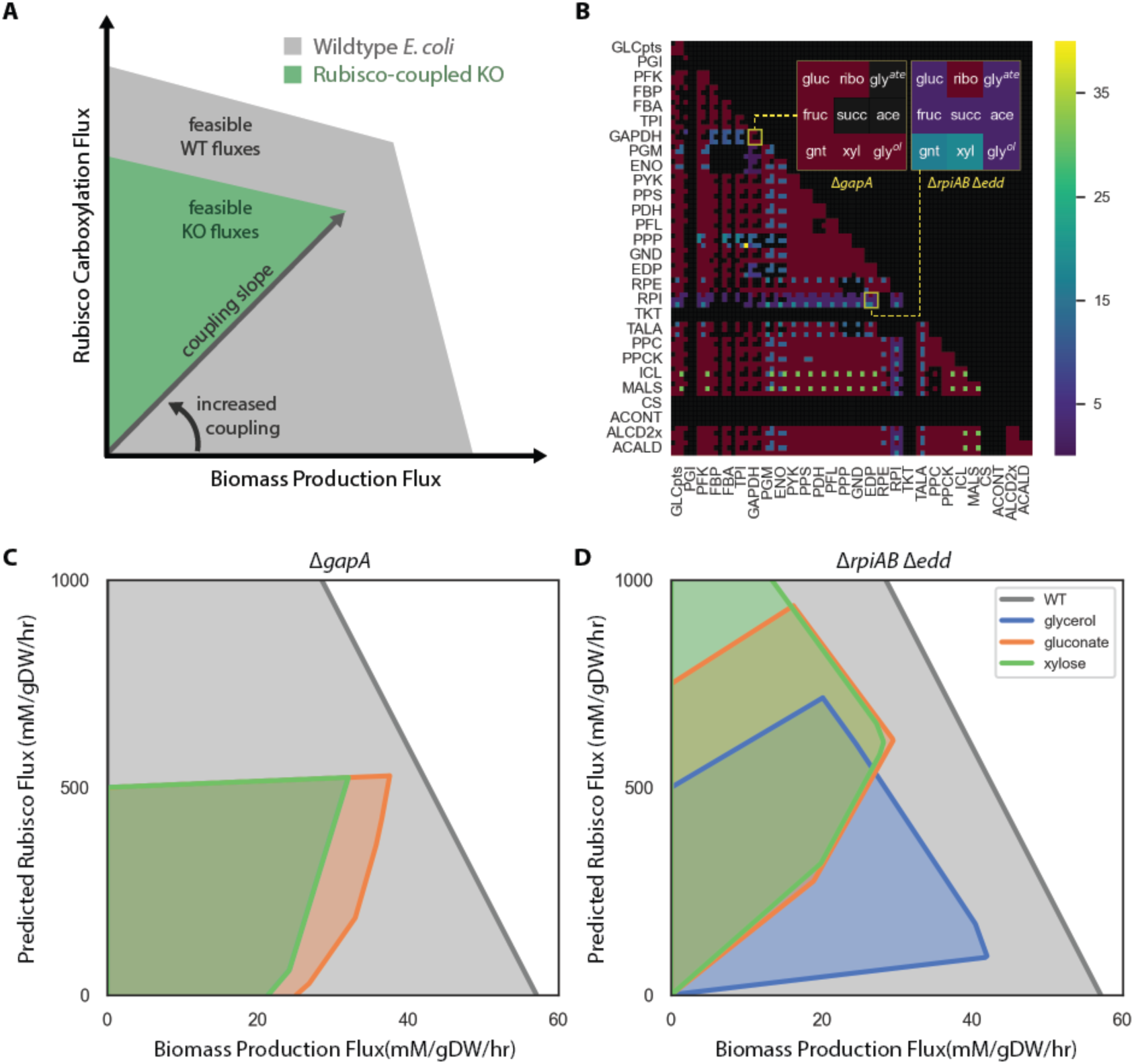
The OptSlope algorithm for designing rubisco-coupled E. coli strains. Optslope searches for metabolic knockout mutants in which biomass production is coupled to flux through a reaction of choice (e.g. rubisco) at all growth rates. (**A**) Shows the space of feasible biomass production and rubisco fluxes for wildtype (WT, grey) and a knockout mutant (green). For WT, biomass production and, therefore, growth rate, are independent of rubisco at all feasible growth rates (i.e. within the grey polygon). The mutant is “rubisco-coupled” because maximal biomass production requires non-zero rubisco carboxylation flux and increasing biomass production demands increased carboxylation. The slope of this relationship is the “coupling slope.” (**B**) We computationally generated pairs of E. coli central metabolic knockouts and calculated the coupling slope on nine carbon sources: glucose (gluc), fructose (fruc), gluconate (gnt), ribose (ribo), succinate (succ), xylose (xyl), glycerate (gly^ate^), acetate (ace) and glycerol (gly^ol^). Each double knockout is summarized as a 3×3 matrix of coupling slopes. Black denotes a rubisco-independent mutant and maroon a coupling slope of 0. The published mutant ΔgapA (Mueller-Cajar et al., 2007) has a coupling slope of 0 (left), while the ΔrpiAB Δedd strain is rubisco-coupled on seven of the carbon sources (right). (**C**) Feasible phase space diagram for the ΔgapA strain shows that biomass production is not coupled to rubisco flux. (**D**) ΔrpiAB Δedd has a positive coupling slope in glycerol, gluconate and xylose media.

**Figure 2 - figure supplement 2.**
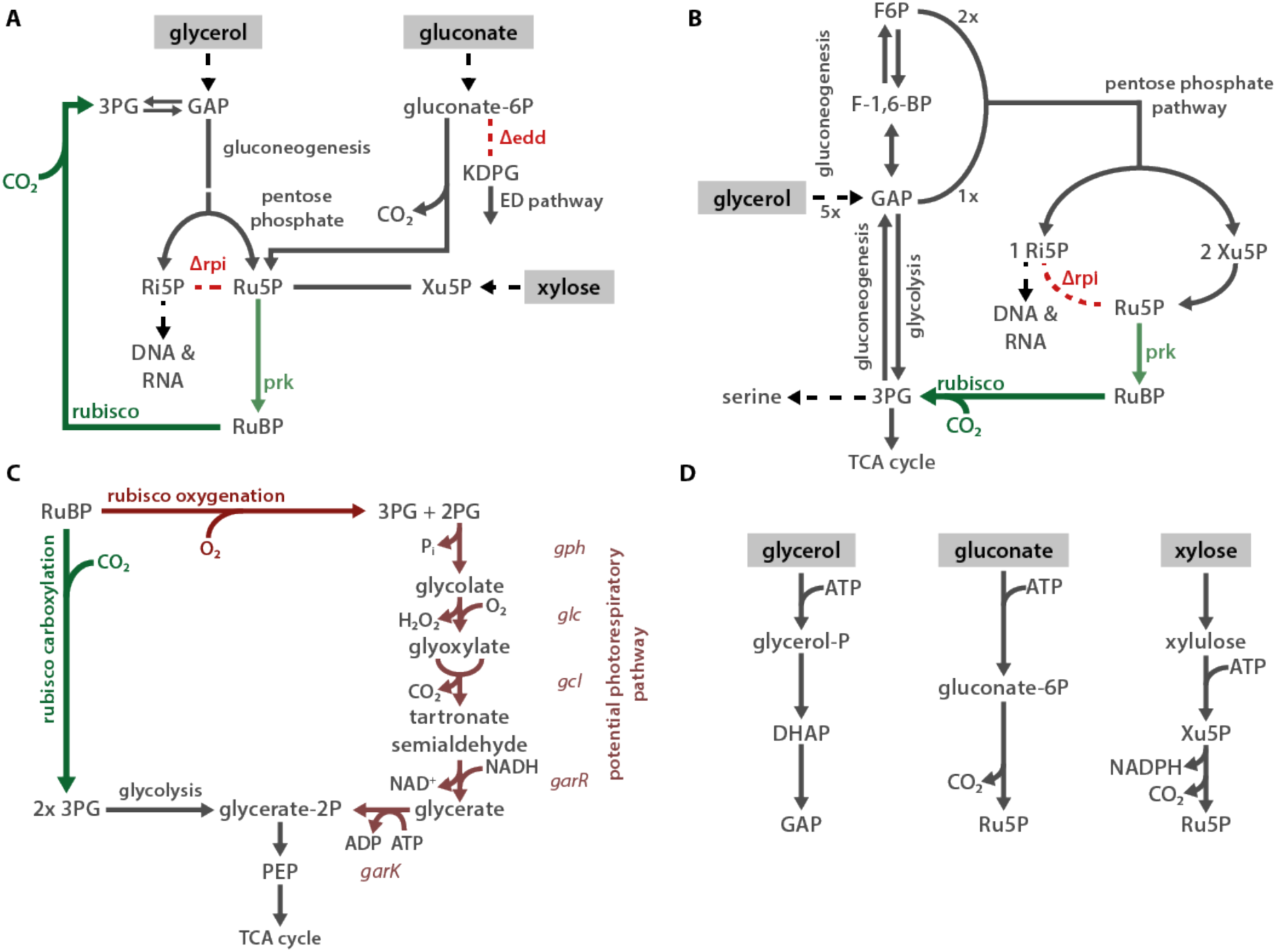
Proposed mechanisms of rubisco-dependent growth in CCMB1. (**A**) CCMB1 depends on rubisco and prk for growth in glycerol, gluconate, and xylose minimal media. The common mechanism is an inability to metabolize ribulose-5-phosphate (Ru5P) due to the deletion of both ribose-phosphate isomerase genes (ΔrpiAB). When gluconate or xylose is the growth substrate, Ru5P must be produced in order to metabolize the carbon source. Though wild type E. coli can metabolize gluconate via the ED pathway, the ED dehydratase knockout (Δedd) in CCMB1 blocks this route and forces 1:1 production of Ru5P from gluconate. Expression of prk and rubisco opens a new route of Ru5P metabolism, thus enabling CCMB1 to grow in gluconate or xylose media. Since extracellular glycerol is converted to glyceraldehyde 3-phosphate (GAP), it can be metabolized through lower glycolysis or through gluconeogenesis. The gluconeogenesis route produces hexoses that enter the pentose phosphate pathway, which is required to synthesize ribose 5-phosphate (Ri5P) for nucleotide and histidine biosynthesis. Depending on the growth rate, products of Ri5P make up 5-25% of E. coli biomass (Bremer and Dennis, 2008; Taymaz-Nikerel et al., 2010). As shown in (**B**), the pentose phosphate pathway forces co-production of Ri5P, Ru5P and xylulose 5-phosphate (Xu5P). In the absence of rpi activity, there is no pathway for metabolism of Xu5P or Ru5P. This defect is complemented by the expression of rubisco and prk. Notably, rubisco can also oxygenate RuBP, as shown in (**C**). E. coli can, in principle, recycle the oxygenation product 2-phosphoglycolate (2PG) though an ersatz photorespiratory pathway via tartronate semialdehyde. This pathway is not the dominant mechanism of rubisco complementation because CCMB1:p1A cannot grow in ambient air, where O_2_ is abundant (Figure 2D). Panel (**D**) describes the initial metabolism of extracellular glycerol, gluconate and xylose in E. coli. Extracellular carbon sources are marked with a grey background throughout. Abbreviations: 3-phosphoglycerate (3PG), 2-phosphoglycolate (2PG), glyceraldehyde 2-phosphate (GAP), dihydroxyacetone phosphate (DHAP), ribose 5-phosphate (Ri5P), ribulose 5-phosphate (Ru5P), xylulose 5-phosphate (Xu5P), ribulose 1,5-bisphosphate (RuBP), 2-keto-3-deoxy-6-phosphogluconate (KDGP), fructose 6-phosphate (F6P), fructose 1,6-bisphosphate (F-1,6-BP), phosphoenolpyruvate (PEP).

**Figure 2 - figure supplement 3.**
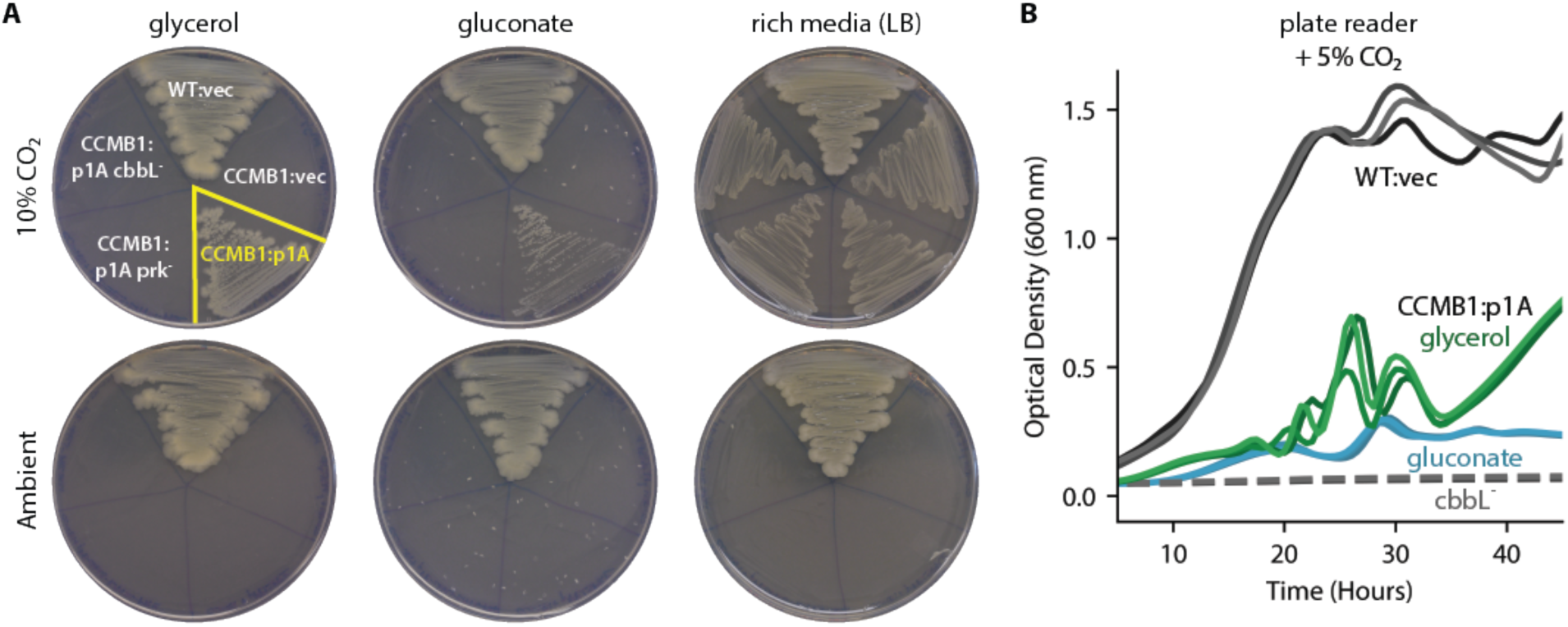
CCMB1 depends on rubisco and prk for growth in minimal media. (**A**) Expression of rubisco and prk complements CCMB1 growth on M9 glycerol and gluconate media under 10% CO_2_, but not in ambient conditions (100 nM aTc induction in M9 plates). Mutations ablating rubisco (cbbL^-^) or prk (prk^-^) activity abrogate growth in selective media but not in LB under 10% CO_2_. Growth in LB is rubisco-independent in 10% CO_2_, but CCMB1 does not grow in ambient air even when supplied rich media because it lacks CA genes (Merlin and Masters, 2003). Growth curves in (**B**) show the rubisco-dependence of CCMB1:p1A growth in glycerol (green) and gluconate (blue) media under 5% CO_2_ in a gas controlled plate reader (Tecan Spark, Methods). Negative controls (CCMB1:p1A cbbL^-^ in glycerol or gluconate media) and uninduced cultures failed to grow in these conditions (dashed grey lines). Though three curves are plotted for each condition in (B), experiments were conducted in technical sextuplicate. Replicates were all consistent.

**Figure 2 - figure supplement 4.**
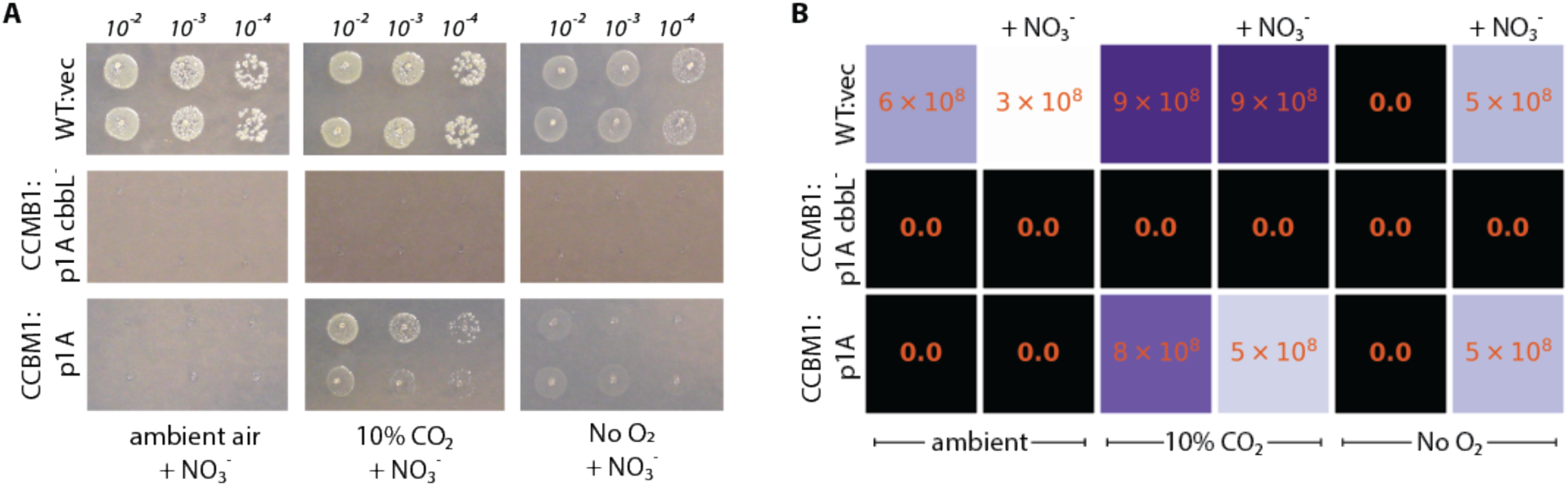
CCMB1 does not require oxygen for growth in minimal media. (**A**) Titer plating assays were used to measure the viability of CCMB1:p1A grown on glycerol media under ambient air (≈0.04% CO_2_, 21% O_2_), 10% CO_2_ (balance air), and an anoxic mix of 10% CO_2_ and 90% N_2_ (“No O_2_”). Since E. coli cannot ferment glycerol, 20 mM NO_3_^-^ was provided as an alternate electron acceptor as marked. (**B**) CCMB1:p1A grows on glycerol media in the absence of O_2_ so long as nitrate is provided. While CCMB1:p1A colonies are noticeably smaller than WT in panel (A), the colony count is indistinguishable, as quantified in panel (B). Experiments were conducted in biological duplicate (i.e. pre-cultures from distinct colonies) with at least two technical replicates (repeated spotting from the same preculture).

**Figure 3 - figure supplement 1.**
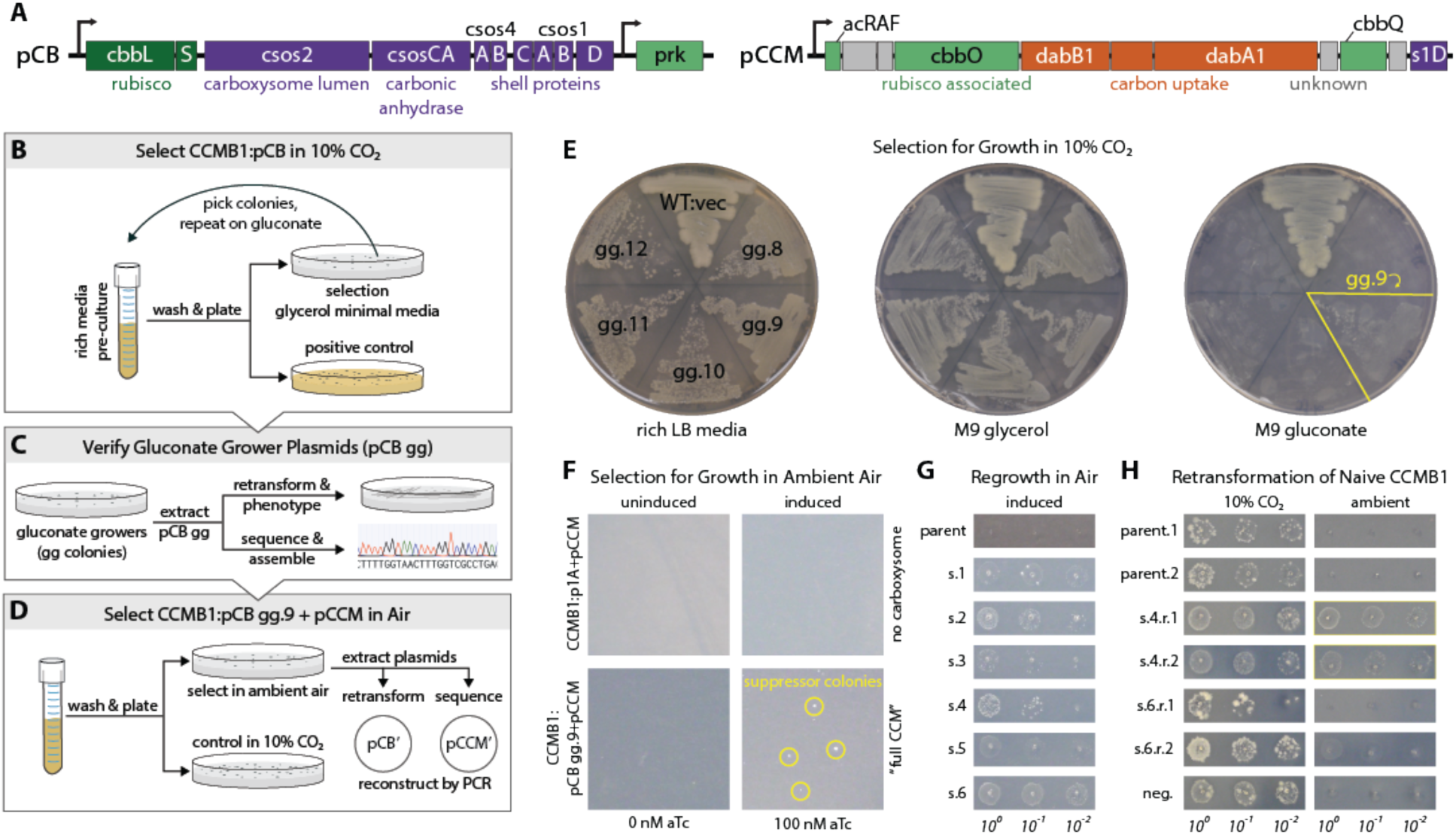
A series of selection experiments produced mutant plasmids that permit rubisco-dependent growth in ambient air. (**A**) pCB and pCCM plasmids together encode 20 H. neapolitanus genes including 12 confirmed CCM components. pCB carries kanamycin resistance and has two transcriptional units expressed under an aTc-inducible P_LtetO-1_ promoter (Lutz and Bujard, 1997). The first derives from pHnCB10 (Bonacci et al., 2012) and expresses 10 carboxysome proteins. The second expresses phosphoribulokinase (prk). pCCM carries chloramphenicol resistance and expresses an 11 gene operon from H. neapolitanus that contains both putative and confirmed CCM genes (Desmarais et al., 2019). Although pCB expresses both rubisco and prk, CCMB1:pCB did not initially grow in M9 media under 10% CO_2_ (not shown) and so we undertook a series of selections, described in panels (B-D) that ultimately led to isolation of pCB’ and pCCM’ plasmids that together enable CCMB1 to grow in ambient air. (**B**) We first selected CCMB1:pCB for growth on minimal media by screening for mutants able to grow on M9 glycerol and then M9 gluconate media. Gluconate growing mutant #9 (gg.9) was used for subsequent experiments as this mutant was found to grow best on gluconate (as shown in E). (**C**) Plasmid extracted from gg.9 was deep sequenced and electroporated into naive CCMB1 to test for plasmid linkage of growth on minimal. (**D**) Selection for rubisco-dependent growth in ambient air. A turbid pre-culture of the CCMB1:pCB gg.9+pCCM double transformant was washed and plated on M9 glycerol media under ambient air. Colonies formed after ≈20 days (as shown in F). 40 colonies (s.1-40) were picked into rich media, grown to saturation, washed and plated on M9 glycerol media to verify growth under ambient air. Roughly 1/4 of chosen colonies regrew under ambient air to varying degrees (s.1-6 are shown in G). Plasmid extracted from several strains was deep-sequenced and electroporated into naive CCMB1 to test plasmid-linkage of growth on glycerol minimal media in ambient air. Pooled plasmid extracted from s.4 was found to confer replicable growth in ambient air (as shown in H). PCR and Gibson cloning were used to reconstruct the individual pCB and pCCM plasmids from this pool. We termed these reconstructed plasmids pCB’ and pCCM’. (**E**) Restreaking of gluconate-growing mutants gg.8-12 described in panel B shows that gg.9 grew best on gluconate. (**F**) CCMB1:pCB gg.9+pCCM double transformants were plated for mutants on M9 glycerol media under ambient air. A negative control lacking carboxysome genes (CCMB1:p1A+pCCM) was plated at the same time. Colonies formed after 20 days (bottom right) only on induced plates (100 nM) and only when all CCM genes were provided (i.e. pCB gg.9 and pCCM). (**G**) Several of the chosen colonies regrew in ambient air. Growth characteristics varied from colony to colony, suggesting genetic variation. (**H**) Pooled plasmid extracted from s.4 was found to permit naive CCMB1 to grow in ambient air. For comparison, plasmid from s.6 produced less reproducible air growth.

**Figure 3 - figure supplement 2.**
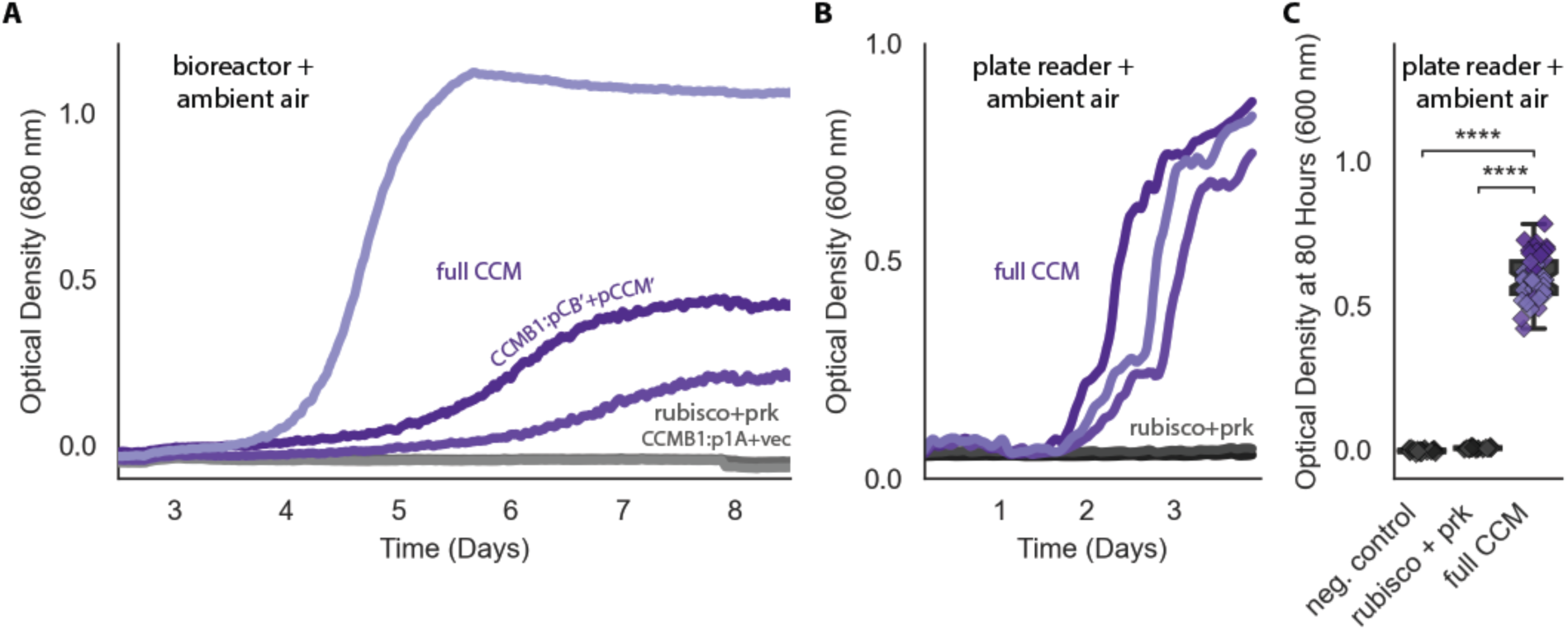
pCB’ and pCCM’ permit CCMB1 to grow in ambient air. (**A**) Biological triplicate growth curves from a bioreactor bubbling ambient air. CCMB1 co-transformed with post-selection plasmids pCB’ and pCCM’ (CCMB1:pCB’ + pCCM’) grows well (purple, “full CCM”), while rubisco and prk alone are insufficient for growth in air (green, “rubisco+prk”). Maximal growth rates for the “full CCM’’ cultures ranged from 0.03-0.06 hr^-1^, corresponding to doubling times of 12-25 hours. As these are biological replicate cultures, heterogeneity in growth kinetics could be due to genetic effects (e.g. point mutations in founding colonies) or non-genetic differences (e.g. varying degree of carboxysome production during pre-culturing). (**B**) Data for the same strains grown in a 96 well plate in ambient air in a shaking plate reader. Different shades mark biological replicates (pre-cultures deriving from three distinct colonies). Additionally, each preculture was used to inoculate at least 12 technical replicates. (**C**) Quantification of the experiment in panel (B) using endpoint data at 80 hours for biological and technical replicates. Panel (C) uses the same colors as (A) and (B) with the addition of a rubisco active site mutant as a negative control (grey, CCMB1:p1A^-^ + vec). ‘****’ indicates P < 10^−10^. P-values were calculated with a Bonferroni-corrected two-sided Mann-Whitney-Wilcoxon test. 10^4^-fold bootstrapping was used to compare “full CCM” data to “rubisco + prk” and estimate a confidence interval for the effect of expressing a full CCM on growth in ambient air, which gave a 99.9% confidence interval of 0.56-0.64 OD units.

**Figure 4 - figure supplement 1.**
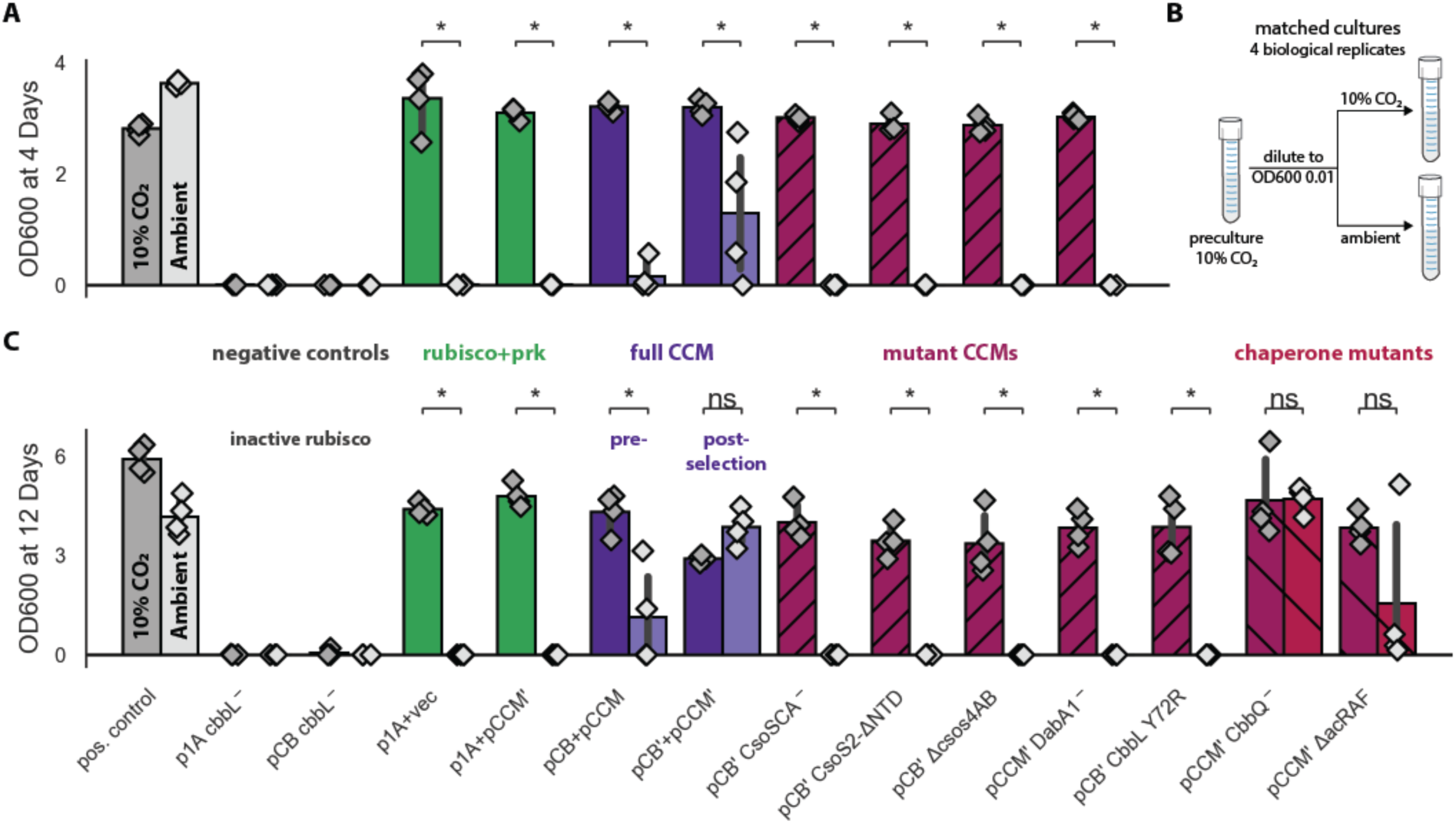
Targeted mutations to the CCM eliminate growth in ambient air. Pre-cultures were grown to saturation in 10% CO_2_ and then diluted to an optical density of 0.01 (600 nm) into two tubes (Methods). One tube was grown in 10% CO_2_ and the other in ambient air, as diagrammed in (**B**). Cells were incubated for 4 days before measuring optical density in (**A**) and 12 days in (**C**). The left bar (darker color) gives the mean endpoint density of biological quadruplicate cultures in 10% CO_2_ and the right bar (lighter color) gives the mean in ambient air. Error bars give a 95% confidence interval of measurements. (A) and (C) share the leftmost 11 strains. From left to right: a positive control (grey, grows in both conditions), two negative controls carrying active site mutants of rubisco (CCMB1:p1A^-^+vec and CCMB1:pCB’^-^+pCCM’), CCMB1 expressing rubisco and prk but no CCM genes (green, CCMB1:p1A+vec) or an incomplete set of CCM genes (green, CCMB1:p1A+pCCM’), CCMB1:pCB+pCCM which carries the pre-selection CCM plasmids (purple), and CCMB1:pCB’+pCCM’ which carries the post-selection plasmids. “vec” denotes an appropriate vector control (pFA-sfGFP). The following pairs of maroon bars describe strains carrying plasmids with targeted CCM mutations: CCMB1:pCB’ CsoSCA^-^+pCCM’ which carries an inactivating mutation to carboxysomal carbonic anhydrase, CCMB1:pCB’ CsoS2 ΔNTD +pCCM’ harboring a deletion of the N-terminal domain of CsoS2 responsible for recruiting rubisco to the carboxysome, CCMB1:pCB’ ΔcsoS4AB + pCCM’ lacking both genes pentameric vertex proteins, and CCMB1:pCB’ DabA1^-^ + pCCM’ carrying an inactivated DAB carbon uptake system. (**A**) CCMB1 grows well in ambient air only when given a full complement of CCM genes on the post-selection plasmids. All mutations to the CCM abrogate growth in air (maroon). Panel (**C**) shows consistent results over a 12-day time period. (C) describes three additional mutants: CCMB1:pCB’ CbbL Y72R + pCCM’ carrying a mutation to the rubisco large subunit that eliminates rubisco-CsoS2 binding, CCMB1:pCB’ + pCCM’ CbbQ^-^ harboring inactivating mutation to the CbbQ subunit of the rubisco activase complex, and CCMB1:pCB’ + pCCM’ ΔacRAF lacking the putative rubisco chaperone acRAF (CCMB1:pCB’ + pCCM’ ΔacRAF). Ablation of rubisco-CsoS2 interaction should eliminate recruitment of rubisco to the carboxyome (Oltrogge et al., 2020). Accordingly, the Y72R mutation eliminated growth in air. Chaperone mutants (CbbQ or acRAF) were both viable in air, though removal of acRAF produced a substantial growth defect (2.5 fold in mean and 8.5 fold in median final density). The positive control strain is the CAfree strain expressing human carbonic anhydrase II (Methods). P-values calculated by a one-sided Mann-Whitney-Wilcoxon test. ‘*’ denotes a P < 0.05. Detailed description of all plasmid abbreviations is given in Table S2.

**Figure 5 - figure supplement 1.**
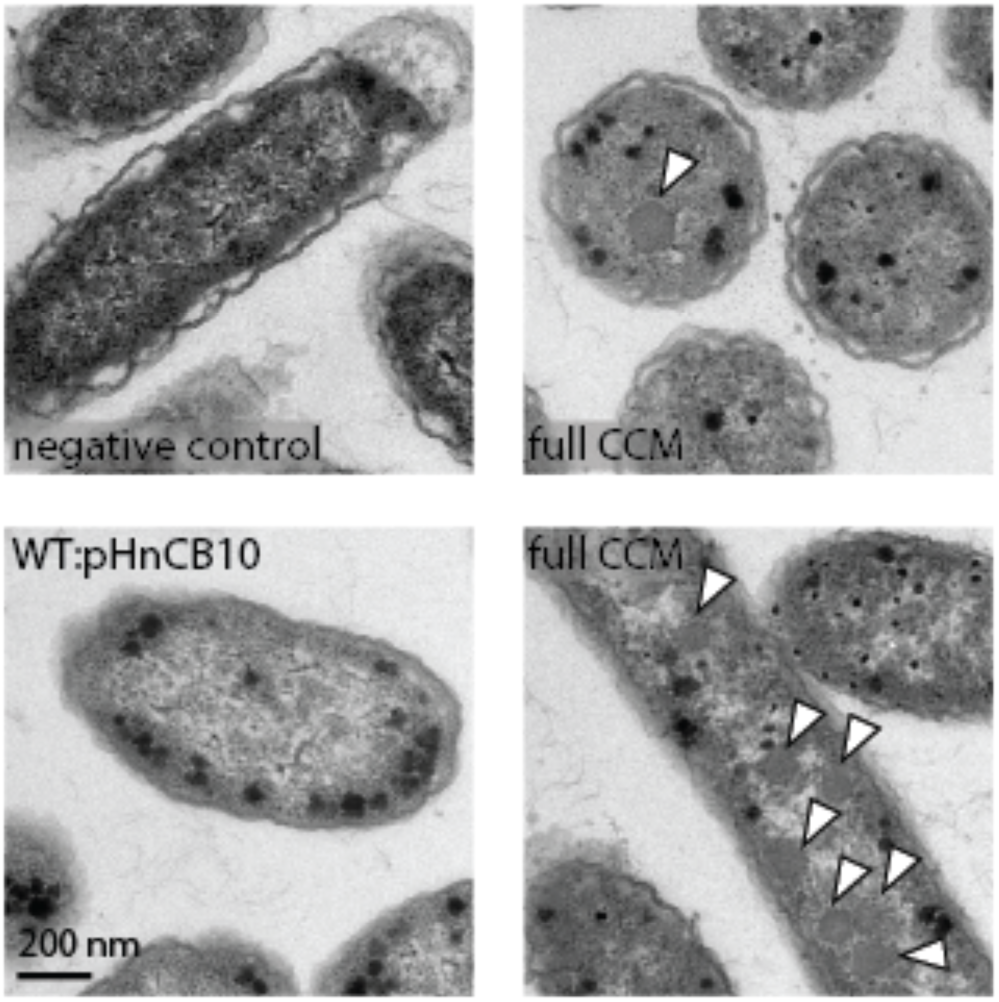
CCMB1:pCB’ + pCCM’ produces polyhedral bodies resembling carboxysomes when grown in ambient air. Transmission electron micrographs of air-grown CCMB1:pCB’+pCCM’ (images on the right) show morphological carboxysomes inside cells (white arrows). The negative control for carboxysome expression is CAfree:pFE-sfGFP + pFA-HCAII (top left). WT:pHnCB10 is the parent strain transformed with a plasmid expressing 10 carboxysome genes and previously shown to enable purification of carboxysome structures from E. coli (Bonacci et al., 2012). This was intended as a positive control, but we did not observe carboxysome structures in electron micrographs of this strain, perhaps because of excessive IPTG induction (500 mM) as previously reported. Expression of carboxysome genes was associated with production of black staining stress granules in both the experiment and pHnCB10 control. These granules were not observed in images of the negative control.

**Figure 5 - figure supplement 2.**
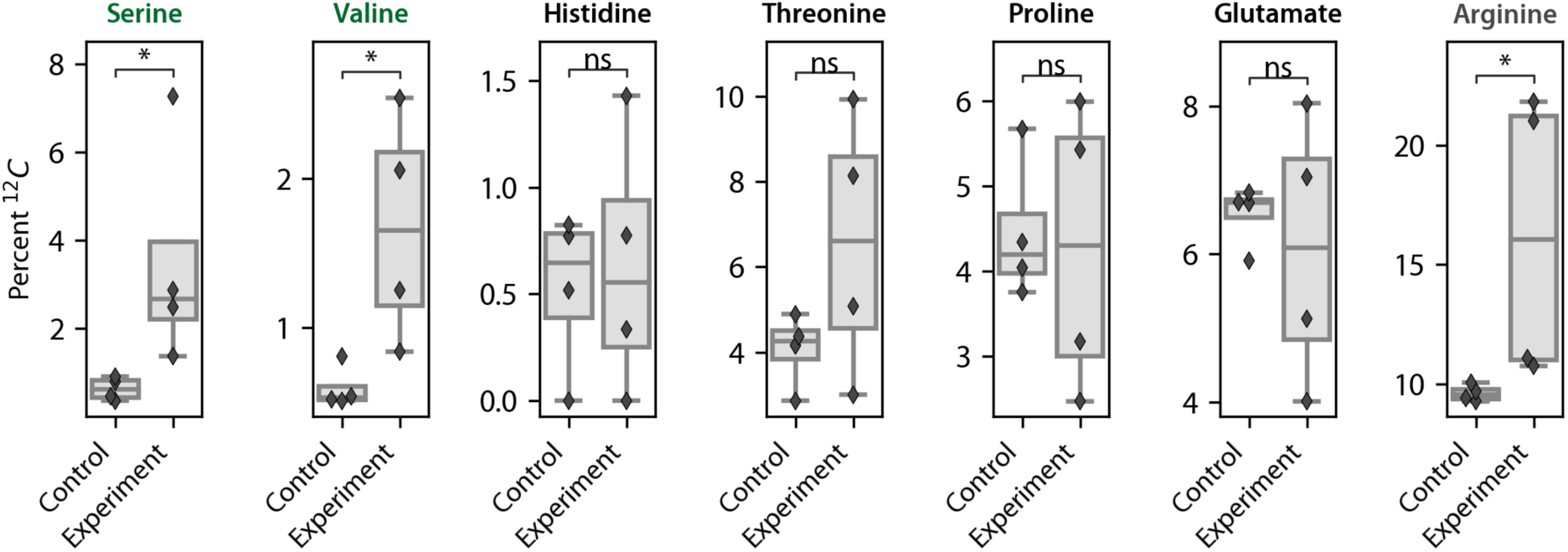
Isotopic composition of amino acids from total biomass hydrolysate. Cells were grown under ambient air in M9 media containing 99% ^13^C labeled glycerol (0.4% v/v) so that nearly all ^12^C in biomass must derive from inorganic carbon. The isotopic composition of amino acids in total biomass hydrolysate of CCMB1:pCB’ + pCCM’ and an appropriate rubisco-independent control were measured via LC-MS (Methods). The control strain is CAfree complemented with the human carbonic anhydrase II, which does not express rubisco (Methods). Serine and valine, which are marked in green, are downstream of the rubisco product 3PG in E. coli central metabolism and, accordingly, show significantly greater ^12^C incorporation in CCMB1:pCB’ + pCCM’ than the control. Histidine, threonine, proline and glutamate are synthesized from precursors deriving from the TCA cycle and pentose phosphate pathways, and thus their carbon atoms do not derive from 3PG (Szyperski, 1995). Arginine is synthesized via a rubisco-independent carboxylation of glutamate (by the addition of carboxyphosphate, (Gleizer et al., 2019)), and so the difference between arginine and glutamate labeling is used to calculate the isotopic composition of intracellular inorganic carbon (C_i_, Methods). Notably, intracellular C_i_ derives both from extracellular C_i_ (predominantly ^12^C) and decarboxylation of the 99% ^13^C glycerol carbon source. As such, the composition will depend on C_i_ uptake as well as the rate of glycerol metabolism. Control cells grew faster than CCMB1:pCB’+pCCM, which can explain why arginine from these cells contains significantly less ^12^C and more ^13^C (from rapid glycerol decarboxylation).

**Figure 5 - figure supplement 1.**
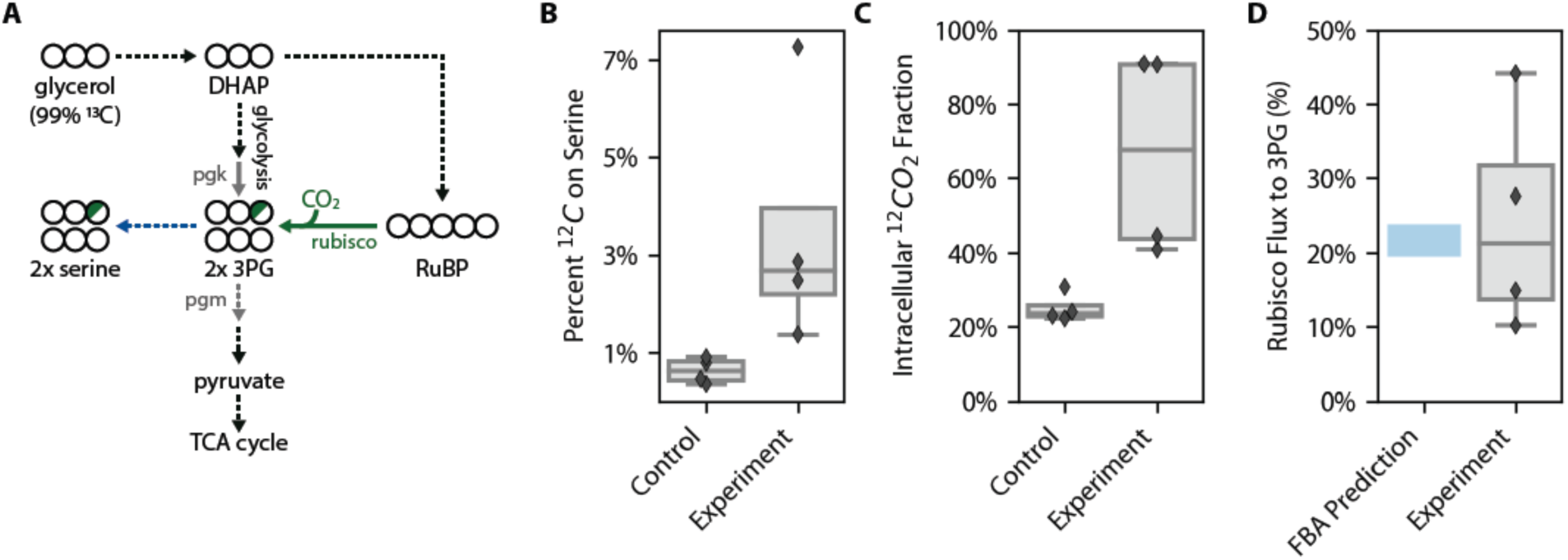
^12^C enrichment on serine is consistent with in vivo CO_2_ fixation. Cells were grown under ambient air in M9 media containing 99% ^13^C labeled glycerol (0.4% v/v) so that nearly all ^12^C in biomass must derive from inorganic carbon. In (**A**) ^13^C atoms are depicted as open circles and fractional ^12^C labeling by a partial green fill color. In CCMB1, 3-phosphoglycerate (3PG) can be produced either through glycolytic metabolism of glycerol (via dihydroxyacetone-phosphate, DHAP) or through rubisco-catalyzed carboxylation of RuBP. At most ⅙ of the carbon atoms on 3PG will be ^12^C when rubisco is active in vivo. In practice this fraction will be less than ⅙ because some of the intracellular inorganic carbon pool (C_i_) derives from decarboxylation of ^13^C labeled glycerol and also because a large fraction of intracellular 3PG is produced through glycolysis (Methods). Serine is a direct metabolic product of 3PG and so reports on the labeling of 3PG. As such, we measured the ^12^C composition of amino acids in total protein hydrolysate via LC-MS (Methods). (**B**) Serine from CCM-expressing CCMB1 cells (‘Experiment’) displayed roughly threefold higher ^12^C labeling than controls, which grow in a rubisco-independent manner (Methods). (**C**) Rubisco carboxylation draws from the intracellular inorganic carbon pool, whose ^12^C composition can be inferred for each sample by comparing the labeling of L-arginine and L-glutamate (Methods). The mean ^12^C fraction of intracellular C_i_ was estimated to be 25% ± 4%and 67% ± 28% for the control and experiment respectively. (**D**) These values were integrated to estimate the percent of 3PG production flux that is due to carboxylation by rubisco (Methods), which was inferred to be 24% ± 15%. These values compare favorably with predictions made via Flux Balance Analysis (19.5-24%, Methods). A sampling method was used to estimate the uncertainty in these rubisco flux inferences (Methods). 99% confidence intervals on the rubisco flux fraction were strictly positive for each biological replicate, with 99% of all posterior estimates between 4% and 51% across all four replicates.

## References

Aguilera J, Van Dijken JP, De Winde JH, Pronk JT. 2005. Carbonic anhydrase (Nce103p): an essential biosynthetic enzyme for growth of Saccharomyces cerevisiae at atmospheric carbon dioxide pressure. Biochem J 391:311–316.

Aigner H, Wilson RH, Bracher A, Calisse L, Bhat JY, Hartl FU, Hayer-Hartl M. 2017. Plant RuBisCo assembly in E. coli with five chloroplast chaperones including BSD2. Science 358:1272–1278.

Andersson I, Knight S, Schneider G, Lindqvist Y, Lundqvist T, Brändén C-I, Lorimer GH. 1989. Crystal structure of the active site of ribulose-bisphosphate carboxylase. Nature 337:229–234.

Antonovsky N, Gleizer S, Noor E, Zohar Y, Herz E, Barenholz U, Zelcbuch L, Amram S, Wides A, Tepper N, Davidi D, Bar-On Y, Bareia T, Wernick DG, Shani I, Malitsky S, Jona G, Bar-Even A, Milo R. 2016. Sugar Synthesis from CO2 in Escherichia coli. Cell 166:115–125.

Baba T, Ara T, Hasegawa M, Takai Y, Okumura Y, Baba M, Datsenko K a., Tomita M, Wanner BL, Mori H. 2006. Construction of Escherichia coli K-12 in-frame, single-gene knockout mutants: the Keio collection. Mol Syst Biol 2:2006.0008.

Bar-Even A, Flamholz A, Noor E, Milo R. 2012. Rethinking glycolysis: on the biochemical logic of metabolic pathways. Nat Chem Biol 8:509–517.

Bar-On YM, Milo R. 2019. The global mass and average rate of rubisco. Proc Natl Acad Sci U S A 116:4738–4743.

Bonacci W, Teng PK, Afonso B, Niederholtmeyer H, Grob P, Silver P a., Savage DF. 2012. Modularity of a carbon-fixing protein organelle. Proc Natl Acad Sci U S A 109:478–483.

Booth IR. 2005. Glycerol and Methylglyoxal Metabolism. EcoSal Plus 1:1–8.

Bowes G, Ogren WL. 1972. Oxygen inhibition and other properties of soybean ribulose 1,5-diphosphate carboxylase. J Biol Chem 247:2171–2176.

Bremer H, Dennis PP. 2008. Modulation of Chemical Composition and Other Parameters of the Cell at Different Exponential Growth Rates. EcoSal Plus 3:1–49.

Cai F, Menon BB, Cannon GC, Curry KJ, Shively JM, Heinhorst S. 2009. The pentameric vertex proteins are necessary for the icosahedral carboxysome shell to function as a CO2 leakage barrier. PLoS One 4:e7521.

Cai Z, Liu G, Zhang J, Li Y. 2014. Development of an activity-directed selection system enabled significant improvement of the carboxylation efficiency of Rubisco. Protein Cell 12–18.

Claassens NJ, Sousa DZ, Dos Santos VAPM, de Vos WM, van der Oost J. 2016. Harnessing the power of microbial autotrophy. Nat Rev Microbiol 14:692–706.

Cleland WW, Andrews TJ, Gutteridge S, Hartman FC, Lorimer GH. 1998. Mechanism of Rubisco: The Carbamate as General Base. Chem Rev 98:549–562.

Datsenko KA, Wanner BL. 2000. One-step inactivation of chromosomal genes in Escherichia coli K-12 using PCR products. Proc Natl Acad Sci U S A 97:6640–6645.

Deatherage DE, Barrick JE. 2014. Identification of mutations in laboratory-evolved microbes from next-generation sequencing data using breseq. Methods Mol Biol 1151:165–188.

Desmarais JJ, Flamholz AI, Blikstad C, Dugan EJ, Laughlin TG, Oltrogge LM, Chen AW, Wetmore K, Diamond S, Wang JY, Savage DF. 2019. DABs are inorganic carbon pumps found throughout prokaryotic phyla. Nat Microbiol 4:2204–2215.

Du J, Förster B, Rourke L, Howitt SM, Price GD. 2014. Characterisation of Cyanobacterial Bicarbonate Transporters in E. coli Shows that SbtA Homologs Are Functional in This Heterologous Expression System. PLoS One 9:e115905.

Ebrahim A, Lerman JA, Palsson BO, Hyduke DR. 2013. COBRApy: COnstraints-Based Reconstruction and Analysis for Python. BMC Syst Biol 7:74.

Eisenhut M, Ruth W, Haimovich M, Bauwe H, Kaplan A, Hagemann M. 2008. The photorespiratory glycolate metabolism is essential for cyanobacteria and might have been conveyed endosymbiontically to plants. Proc Natl Acad Sci U S A 105:17199–17204.

Ermakova M, Danila FR, Furbank RT, von Caemmerer S. 2020. On the road to C4 rice: advances and perspectives. Plant J 101:940–950.

Fischer WW, Hemp J, Johnson JE. 2016. Evolution of Oxygenic Photosynthesis. Annu Rev Earth Planet Sci 44:647–683.

Flamholz AI, Prywes N, Moran U, Davidi D, Bar-On YM, Oltrogge LM, Alves R, Savage D, Milo R. 2019. Revisiting Trade-offs between Rubisco Kinetic Parameters. Biochemistry 58:3365–3376.

Flamholz A, Shih PM. 2020. Cell biology of photosynthesis over geologic time. Curr Biol 30:R490–R494.

Gleizer S, Ben-Nissan R, Bar-On YM, Antonovsky N, Noor E, Zohar Y, Jona G, Krieger E, Shamshoum M, Bar-Even A, Milo R. 2019. Conversion of Escherichia coli to Generate All Biomass Carbon from CO2. Cell 179:1255–1263.e12.

Herz E, Antonovsky N, Bar-On Y, Davidi D, Gleizer S, Prywes N, Noda-Garcia L, Frisch KL, Zohar Y, Wernick DG, Others. 2017. The genetic basis for the adaptation of E. coli to sugar synthesis from CO 2. Nat Commun 8:1705.

Higgins CF, Hiles ID, Salmond GP, Gill DR, Downie JA, Evans IJ, Holland IB, Gray L, Buckel SD, Bell AW. 1986. A family of related ATP-binding subunits coupled to many distinct biological processes in bacteria. Nature 323:448–450.

Holzhütter H-G. 2004. The principle of flux minimization and its application to estimate stationary fluxes in metabolic networks. Eur J Biochem 271:2905–2922.

Iñiguez C, Capó-Bauçà S, Niinemets Ü, Stoll H, Aguiló-Nicolau P, Galmés J. 2020. Evolutionary trends in RuBisCO kinetics and their co-evolution with CO2 concentrating mechanisms. Plant J 101:897–918.

Kerfeld CA, Melnicki MR. 2016. Assembly, function and evolution of cyanobacterial carboxysomes. Curr Opin Plant Biol 31:66–75.

Lewis NE, Nagarajan H, Palsson BO. 2012. Constraining the metabolic genotype–phenotype relationship using a phylogeny of in silico methods. Nat Rev Microbiol 10:291–305.

Liang S, Bipatnath M, Xu Y, Chen S, Dennis P, Ehrenberg M, Bremer H. 1999. Activities of constitutive promoters in Escherichia coli. J Mol Biol 292:19–37.

Lin MT, Occhialini A, Andralojc PJ, Parry MAJ, Hanson MR. 2014. A faster Rubisco with potential to increase photosynthesis in crops. Nature 513:547–550.

Long BM, Hee WY, Sharwood RE, Rae BD, Kaines S, Lim Y-L, Nguyen ND, Massey B, Bala S, von Caemmerer S, Badger MR, Price GD. 2018. Carboxysome encapsulation of the CO2-fixing enzyme Rubisco in tobacco chloroplasts. Nat Commun 9:3570.

Long BM, Rae BD, Rolland V, Förster B, Price GD. 2016. Cyanobacterial CO2-concentrating mechanism components: function and prospects for plant metabolic engineering. Curr Opin Plant Biol 31:1–8.

Lutz R, Bujard H. 1997. Independent and tight regulation of transcriptional units in Escherichia coli via the LacR/O, the TetR/O and AraC/I1-I2 regulatory elements. Nucleic Acids Res 25:1203–1210.

Mackinder LCM, Meyer MT, Mettler-Altmann T, Chen VK, Mitchell MC, Caspari O, Freeman Rosenzweig ES, Pallesen L, Reeves G, Itakura A, Roth R, Sommer F, Geimer S, Mühlhaus T, Schroda M, Goodenough U, Stitt M, Griffiths H, Jonikas MC. 2016. A repeat protein links Rubisco to form the eukaryotic carbon-concentrating organelle. Proc Natl Acad Sci U S A 113:5958–5963.

Mangan NM, Flamholz A, Hood RD, Milo R, Savage DF. 2016. pH determines the energetic efficiency of the cyanobacterial CO2 concentrating mechanism. Proc Natl Acad Sci U S A 113:E5354–62.

Marcus Y, Schwarz R, Friedberg D, Kaplan A. 1986. High CO2 Requiring Mutant of Anacystis nidulans R2. Plant Physiol 82:610–612.

McGrath JM, Long SP. 2014. Can the cyanobacterial carbon-concentrating mechanism increase photosynthesis in crop species? A theoretical analysis. Plant Physiol 164:2247–2261.

Merlin C, Masters M. 2003. Why is carbonic anhydrase essential to Escherichia coli? J Bacteriol 185. doi:10.1128/JB.185.21.6415

Mueller-Cajar O. 2017. The Diverse AAA+ Machines that Repair Inhibited Rubisco Active Sites. Front Mol Biosci 4:31.

Mueller-Cajar O, Morell M, Whitney SM. 2007. Directed evolution of rubisco in Escherichia coli reveals a specificity-determining hydrogen bond in the form II enzyme. Biochemistry 46:14067–14074.

Nevins CP, Vierck JL, Bogachus LD, Velotta NS, Castro-Munozledo F, Dodson MV. 2005. An Inexpensive Method for Applying Nitrogen Evaporation to Hexane-containing 24- or 96-well Plates. Cytotechnology 49:71–75.

Occhialini A, Lin MT, Andralojc PJ, Hanson MR, Parry MAJ. 2016. Transgenic tobacco plants with improved cyanobacterial Rubisco expression but no extra assembly factors grow at near wild-type rates if provided with elevated CO2. Plant J 85:148–160.

Oltrogge LM, Chaijarasphong T, Chen AW, Bolin ER, Marqusee S, Savage DF. 2020. Multivalent interactions between CsoS2 and Rubisco mediate α-carboxysome formation. Nat Struct Mol Biol 27:281–287.

Orr DJ, Worrall D, Lin MT, Carmo-Silva E, Hanson MR, Parry MAJ. 2020. Hybrid Cyanobacterial-Tobacco Rubisco Supports Autotrophic Growth and Procarboxysomal Aggregation. Plant Physiol 182:807–818.

Orth JD, Fleming RMT, Palsson BØ. 2010. Reconstruction and Use of Microbial Metabolic Networks: the Core Escherichia coli Metabolic Model as an Educational Guide. EcoSal Plus 4:1–47.

Peekhaus N, Conway T. 1998. What’s for dinner : Entner-Doudoroff metabolism in Escherichia coli. J Bacteriol 180:3495.

Pellicer MT, Nuñez MF, Aguilar J, Badia J, Baldoma L. 2003. Role of 2-Phosphoglycolate Phosphatase of Escherichia coli in Metabolism of the 2-Phosphoglycolate Formed in DNA Repair. J Bacteriol 185:5815–5821.

Price GD, Badger MR. 1989a. Isolation and characterization of high CO2-requiring-mutants of the cyanobacterium Synechococcus PCC7942: two phenotypes that accumulate inorganic carbon but are apparently unable to generate CO2 within the carboxysome. Plant Physiol 91:514–525.

Price GD, Badger MR. 1989b. Expression of Human Carbonic Anhydrase in the Cyanobacterium Synechococcus PCC7942 Creates a High CO2-Requiring Phenotype Evidence for a Central Role for Carboxysomes in the CO2 Concentrating Mechanism. Plant Physiol 91:505–513.

Rae BD, Long BM, Badger MR, Price GD. 2013. Functions, compositions, and evolution of the two types of carboxysomes: polyhedral microcompartments that facilitate CO2 fixation in cyanobacteria and some proteobacteria. Microbiol Mol Biol Rev 77:357–379.

Raven JA, Beardall J, Sánchez-Baracaldo P. 2017. The possible evolution and future of CO2-concentrating mechanisms. J Exp Bot 68:3701–3716.

Sage RF, Sage TL, Kocacinar F. 2012. Photorespiration and the evolution of C4 photosynthesis. Annu Rev Plant Biol 63:19–47.

Sawaya MR, Cannon GC, Heinhorst S, Tanaka S, Williams EB, Yeates TO, Kerfeld C a. 2006. The structure of beta-carbonic anhydrase from the carboxysomal shell reveals a distinct subclass with one active site for the price of two. J Biol Chem 281:7546–7555.

Scott KM, Leonard JM, Boden R, Chaput D, Dennison C, Haller E, Harmer TL, Anderson A, Arnold T, Budenstein S, Brown R, Brand J, Byers J, Calarco J, Campbell T, Carter E, Chase M, Cole M, Dwyer D, Grasham J, Hanni C, Hazle A, Johnson C, Johnson R, Kirby B, Lewis K, Neumann B, Nguyen T, Nino Charari J, Morakinyo O, Olsson B, Roundtree S, Skjerve E, Ubaldini A, Whittaker R. 2019. Diversity in CO2-Concentrating Mechanisms among Chemolithoautotrophs from the Genera Hydrogenovibrio, Thiomicrorhabdus, and Thiomicrospira, Ubiquitous in Sulfidic Habitats Worldwide. Appl Environ Microbiol 85:1–19.

Sezonov G, Joseleau-Petit D, D’Ari R. 2007. Escherichia coli physiology in Luria-Bertani broth. J Bacteriol 189:8746–8749.

Shih PM, Occhialini A, Cameron JC, Andralojc PJ, Parry MAJ, Kerfeld CA. 2016. Biochemical characterization of predicted Precambrian RuBisCO. Nat Commun 7:10382.

Stauffer GV. 2004. Regulation of Serine, Glycine, and One-Carbon Biosynthesis. EcoSal Plus 1:1–22.

Stolper DA, Revsbech NP, Canfield DE. 2010. Aerobic growth at nanomolar oxygen concentrations. Proc Natl Acad Sci U S A 107:18755–18760.

Szyperski T. 1995. Biosynthetically directed fractional 13C-labeling of proteinogenic amino acids. An efficient analytical tool to investigate intermediary metabolism. Eur J Biochem 232:433–448.

Taymaz-Nikerel H, Borujeni AE, Verheijen PJT, Heijnen JJ, van Gulik WM. 2010. Genome-derived minimal metabolic models for Escherichia coli MG1655 with estimated in vivo respiratory ATP stoichiometry. Biotechnol Bioeng 107:369–381.

Tsai Y-CC, Lapina MC, Bhushan S, Mueller-Cajar O. 2015. Identification and characterization of multiple rubisco activases in chemoautotrophic bacteria. Nat Commun 6:8883.

Unden G, Dünnwald P. 2008. The Aerobic and Anaerobic Respiratory Chain of Escherichia coli and Salmonella enterica: Enzymes and Energetics. EcoSal Plus 3. doi:10.1128/ecosalplus.3.2.2

Wang L, Jonikas MC. 2020. The pyrenoid. Curr Biol 30:R456–R458.

Wheatley NM, Sundberg CD, Gidaniyan SD, Cascio D, Yeates TO. 2014. Structure and identification of a pterin dehydratase-like protein as a ribulose-bisphosphate carboxylase/oxygenase (RuBisCO) assembly factor in the α-carboxysome. J Biol Chem 289:7973–7981.

Wildman SG. 2002. Along the trail from Fraction I protein to Rubisco (ribulose bisphosphate carboxylase-oxygenase). Photosynth Res 73:243–250.

Wilson RH, Martin-Avila E, Conlan C, Whitney SM. 2018. An improved Escherichia coli screen for Rubisco identifies a protein-protein interface that can enhance CO2-fixation kinetics. J Biol Chem 293:18–27.

Winkler ME, Ramos-Montañez S. 2009. Biosynthesis of Histidine. EcoSal Plus 3:1–33.

Wu A, Hammer GL, Doherty A, von Caemmerer S, Farquhar GD. 2019. Quantifying impacts of enhancing photosynthesis on crop yield. Nat Plants 5:380–388.

